# Prospective discovery of LasB inhibitors by integrating generative design, free energy ranking, and experimental validation

**DOI:** 10.64898/2026.10.12.758448

**Authors:** Alejandro Martínez León, Angeliki Vgenopoulou, Gwenaëlle Jézéquel, Andreas M. Kany, Anna K. H. Hirsch, Jochen S. Hub

## Abstract

Identifying synthetically accessible, potent ligands from vast chemical space remains a major challenge in drug discovery, particularly because experimental synthesis and testing are costly and low throughput. Here, we present a fully open-source computational–experimental workflow that integrates fragment-based *de novo* molecular design with rigorous alchemical absolute binding free-energy calculations to enable physics-based prioritization before chemical synthesis. We apply this strategy by targeting the virulence-associated zinc metalloprotease LasB from *Pseudomonas aeruginosa*. From ∼40,000 diverse compounds generated through multi-objective optimization, a small subset was prioritized and experimentally evaluated, leading to the discovery of three previously unreported nanomolar inhibitors (IC_50_ 22–53 nM) with excellent metabolic and plasma stability. This work demonstrates a general and scalable framework for coupling generative molecular design with quantitative binding affinity prediction, improving decision-making and reducing the cost, time, and risk associated with early-stage drug discovery.

**Graphical abstract:** 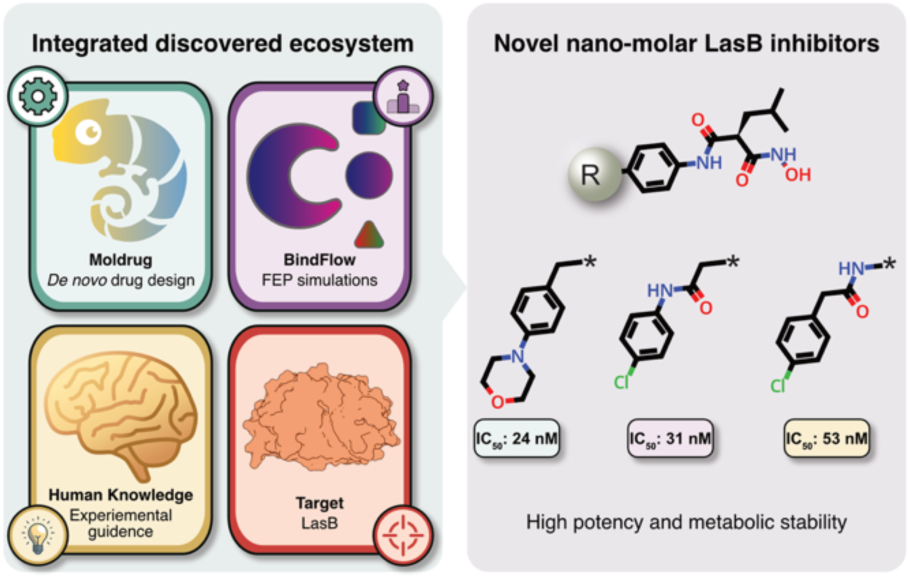

## Introduction

Drug discovery and development is a costly and time-consuming process^1,2^, and computational methods have become essential for accelerating and streamlining many of its stages, including hit identification, molecular optimization, and binding-affinity prediction^3^. Virtual screening methods^3,4^ have become central to hit identification, while *de novo* drug design algorithms based on machine learning^5,6^, reaction rules^7^, or fragment-based strategies^8–10^ play an increasingly important role in molecular optimization and the generation of novel candidate molecules. For binding-affinity prediction, alchemical binding free energy methods, commonly referred to as free energy perturbation (FEP), represent one of the most rigorous and physically grounded approaches available. FEP relies on statistical thermodynamics to estimate binding free energies and has demonstrated remarkable accuracy, often achieving root mean squared errors (RMSE) of 1−2 kcal mol^−1^ relative to experimental measurements^11,12^, thus approaching experimental precision^13^.

*De novo* drug design has demonstrated strong potential for the exploration of chemical space^14^, whereas FEP has proven particularly powerful as a post-ranking tool to prioritize binders for synthesis and experimental validation via relative (RBFE) or absolute binding free energy (ABFE) calculations^15–21^. While *de novo* drug design and FEP are complementary, their combined prospective application remains underrepresented in the literature, with only a few notable examples reported to date^21–28^. In addition, most reported studies preferentially employ RBFE methodologies, with only a limited number implementing ABFE calculations^26,27^. In our view, the limited adoption of integrated *de novo* design and FEP workflows largely stems from their steep learning curves and the high costs associated with commercial software licenses.

Recently, we addressed these challenges by developing two validated, user-friendly, and open-source tools: Moldrug^29^, a fragment-based generative and multi-objective optimization platform for *de novo* molecular design, and BindFlow^12^, an end-to-end automated workflow for performing ABFE calculations at scale. Together, these tools offer a framework that spans from ideation to physics-based prioritization. Here, we demonstrate their use in tandem by applying them to a real-world challenge: the design of new compounds targeting *Pseudomonas aeruginosa* LasB.

*P. aeruginosa* is a Gram-negative bacterium with increasing global prevalence of multi-drug resistant strains.^30^ *P. aeruginosa* infections are a leading cause of hospital-acquired mortality, which were associated with approximately 560,000 deaths globally in 2019.^31^ *P. aeruginosa* is challenging to treat due to its intrinsic resistance mechanisms, its ability to acquire additional resistance determinants, and its extensive repertoire of virulence factors.^32^ Among these virulence factors, pseudolysin (also known as elastase B or LasB, named after its encoding gene)^33,34^ is the most predominant extracellular virulence factor and plays a central role in pathogenesis.^35,36^ LasB is a secreted zinc-metalloprotease that degrades components of the extracellular matrix, disrupts epithelial and endothelial barriers, and interferes with host immune responses^35^ (Figure 1). Through these activities, LasB facilitates nutrient acquisition, promotes bacterial dissemination, immune evasion and contributes to tissue damage and poor clinical outcomes.^35^

**Figure 1.**
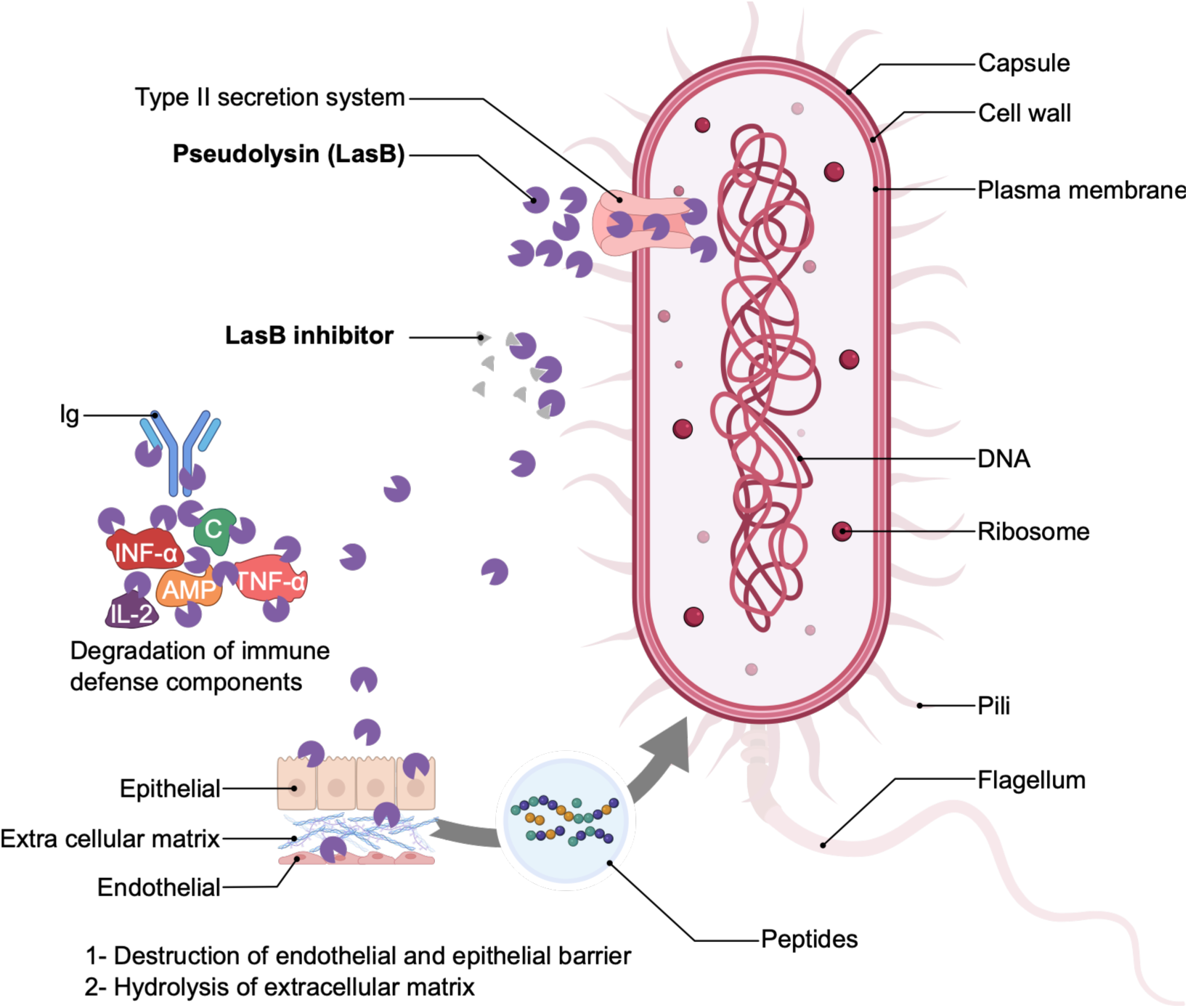
Pseudomonas aeruginosa. Biological role of LasB. Diagram inspired by Ref. 34. For explanation, see text.

Targeting LasB presents several conceptual advantages for therapeutic development.^36^ First, because LasB is secreted into the extracellular space, inhibitors do not need to cross the complex outer membrane of *P. aeruginosa* and accumulate inside the cell, circumventing a key hurdle in anti-Gram-negative drug discovery. Second, antivirulence strategies aimed at neutralizing pathogenic mechanisms—rather than killing the bacterium—are expected to impose lower evolutionary pressure, thereby reducing the likelihood of resistance development.^35,37^ Third, selective inhibition of LasB could attenuate disease severity^38–41^ while preserving the commensal microbiota and minimizing off-target effects associated with broad-spectrum antibiotics^42,43^. LasB contains a conserved *Zn*²⁺-binding catalytic site, and several inhibitor classes have been described, most of which involve coordination of the metal center.^44–53^

In previous works^53,54^, we determined the X-ray cocrystal structure of LasB in complex with a hydroxamic acid inhibitor, revealing a bidentate coordination where both the carbonyl and hydroxamate oxygen atom interact with the active site *Zn*^2+^ ion (Figure 2b). This interaction results in a distorted trigonal bipyramidal geometry, further stabilized by a network of polar residues, including His140, Glu141, and the backbone carbonyl of Ala113. In addition, the isobutyl substituent of molecules **2** and **3** (Figure 2c) effectively occupies a hydrophobic subpocket contributing to the overall affinity.^53^

**Figure 2.**
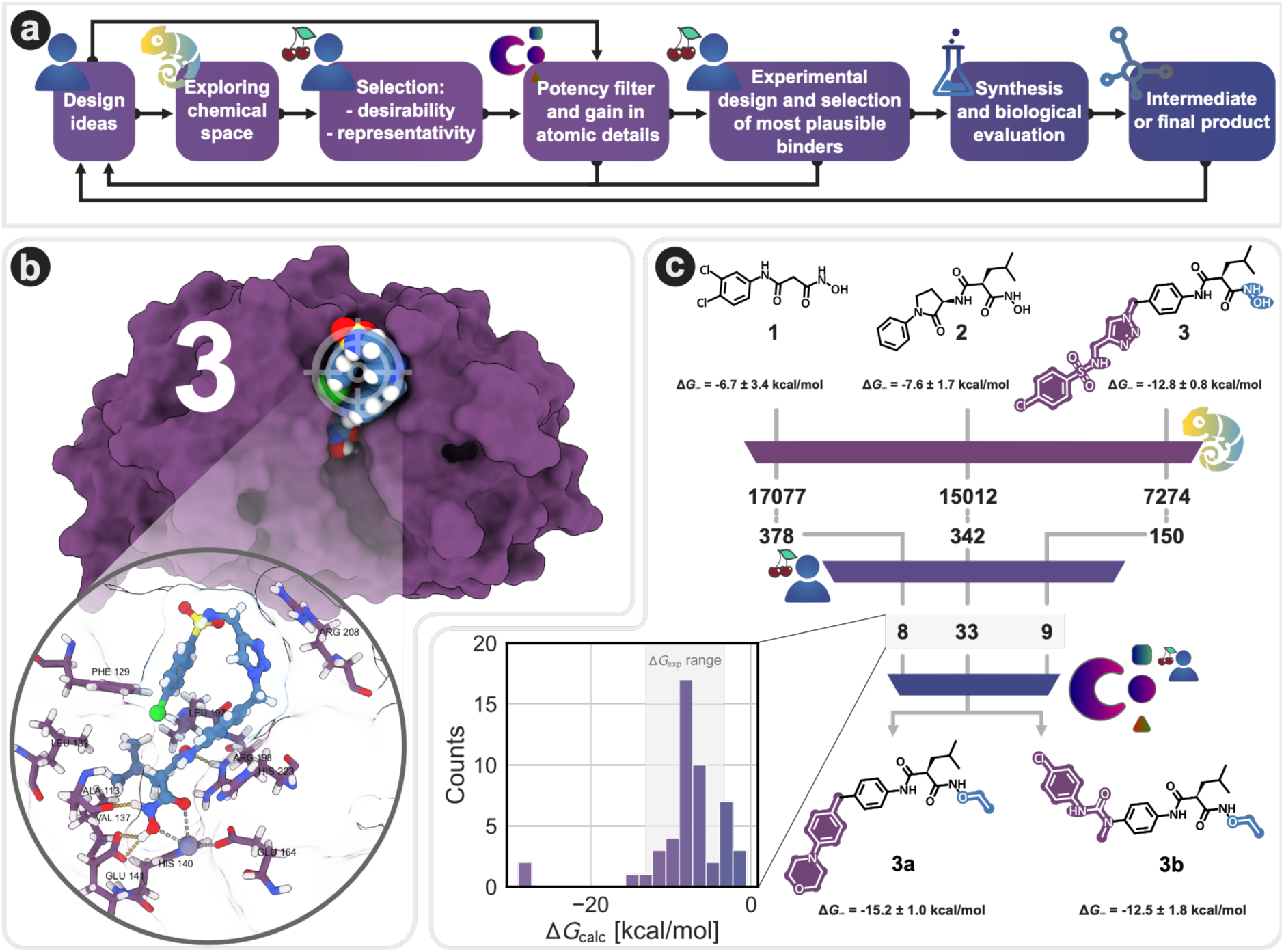
Iterative workffow for structure-based drug design and optimization. **(a)** Overview of the design workffow. **(b)** Seed molecule **3** in complex with LasB (PDB: 32KK^57^); a close-up view of the binding pose highlights key interacting residues. Zn^2+^ interactions are shown as light-gray, dashed lines, and hydrogen bonds as yellow, dashed lines. **(c)** Chemical structures of the seed molecules (**1**-**3** ^53,57–59^) and selected de novo designed molecules (**3a** and **3b**), along with their estimated binding free energies. The computational workffow is illustrated, starting with Moldrug-based design, followed by visual inspection and manual selection of 50 chemically diverse molecules. Final candidate selection is based on BindFlow predictions and practical considerations. The histogram shows the distribution of estimated binding free energies for the 50 selected molecules, with the experimental range of the HIPS internal molecules indicated by light-gray shading.

Although our group has synthesized a variety of hydroxamic acid-based inhibitors, with low nanomolar IC50 values and high selectivity over several tested human matrix metalloproteases^55,56^, previous structure–activity relationship (SAR) studies have focused heavily on the zinc-binding group and the immediate hydrophobic subpocket. Consequently, the chemical space and potential interactions in the regions surrounding the active site remain unexplored.

In this study, we explored the LasB binding landscape through fragment-based multi-objective optimization^29^, generating 870 candidate molecules within 39,364 structures sampled during the process. From this pool, a chemically diverse subset of 50 compounds was prioritized based on predicted drug-like relevant scores and structural diversity. These compounds were then subjected to binding ABFE calculations^12^ to prioritize for synthesis, leading to the experimental validation of six novel compounds, three of which demonstrated nanomolar affinity for LasB. While these binders showed excellent metabolic and plasma stability, some cytotoxicity was noted. Ultimately, this workflow illustrates the power of integrating *de novo* design, FEP-based free-energy ranking, and experimental expertise to efficiently triage vast chemical spaces and drive informed decision-making in drug discovery.

## Results

To systematically discover and optimize novel LasB inhibitors, we implemented an integrated, iterative structure-based drug design workflow (Figure 2a). This pipeline conceptually bridges computational generation with experimental validation. The process begins with the exploration of chemical space starting from initial design ideas, followed by selection criteria based on desirability profiles and structural diversity. Generated compounds then pass through a rigorous potency filter before the most plausible binders are prioritized for synthesis and biological evaluation.

### Overview of the design process

Figure 2c illustrates the application of this general workflow within the current study. We started from three seed molecules (**1**-**3**), for which the protein–ligand complex and binding affinity were known^53,57–59^. We carried out seven design campaigns, each performed in triplicate (see *Methods*). The Moldrug-driven design took as input those three seed molecules and was guided by consensus scores constructed by: Vina docking score^60^, Quantitative Estimation of Drug-likeness (QED)^61^ and Synthetic Accessibility (SA)^62^ score. This strategy ensured a balance between predicted potency, drug-likeness, and synthetic accessibility.

Our exploration focused primarily on modifications of regions not associated with *Zn*^2+^-binding for three reasons: (i) the well-established importance of the hydroxamic acid moiety for *Zn*^2+^ coordination, (ii) force field uncertainties owing to *Zn*^2+^ interactions during MD simulations, and (iii) the limited prior exploration of the non-*Zn*^2+^-binding region.

Nevertheless, we also explored molecular growth in the direction of the hydroxamic acid hydroxyl group while introducing only minimal changes to the *Zn*^2+^-binding region of the seed molecules. Specifically, campaigns I, II, and VII allowed molecular growth via the hydroxyl hydrogen, whereas campaign V focused exclusively on growth through this hydroxyl group (see *Methods*).

While conventional virtual screening typically requires the brute-force evaluation of libraries comprising millions or billions of molecules^4,16,63^, Moldrug achieves an extensive exploration of chemical space requiring only a few thousand generated candidates to identify optimized solutions.^29^ In this study, Moldrug navigated the chemical landscape by generating only 39,364 molecules (Figure 2c). To evaluate the range and diversity of this exploration, we performed a t-SNE analysis to visualize the chemical-space coverage.

As shown in the t-SNE representation (Figure 3a), Moldrug-generated molecules define the broad boundaries of the explored chemical space. Conversely, the internal, experimental HIPS dataset of elastase inhibitors (Figure 3a, green dots), which was accumulated over years of experimental work, occupies only a small, central region. Notably, the search originated from only three localized seed molecules (Figure 3a, brown dots), representing a narrow starting point within the known chemical landscape. These observations indicate that Moldrug effectively drives exploration toward underrepresented and previously unexplored regions of the chemical space, meaning the final chemical coverage is neither restricted nor biased by the initial positioning of the seeds. Ultimately, we demonstrated that a minimal starting point is sufficient for Moldrug to conduct a diverse and exhaustive exploration, which aligns with previous studies.^29^

**Figure 3.**
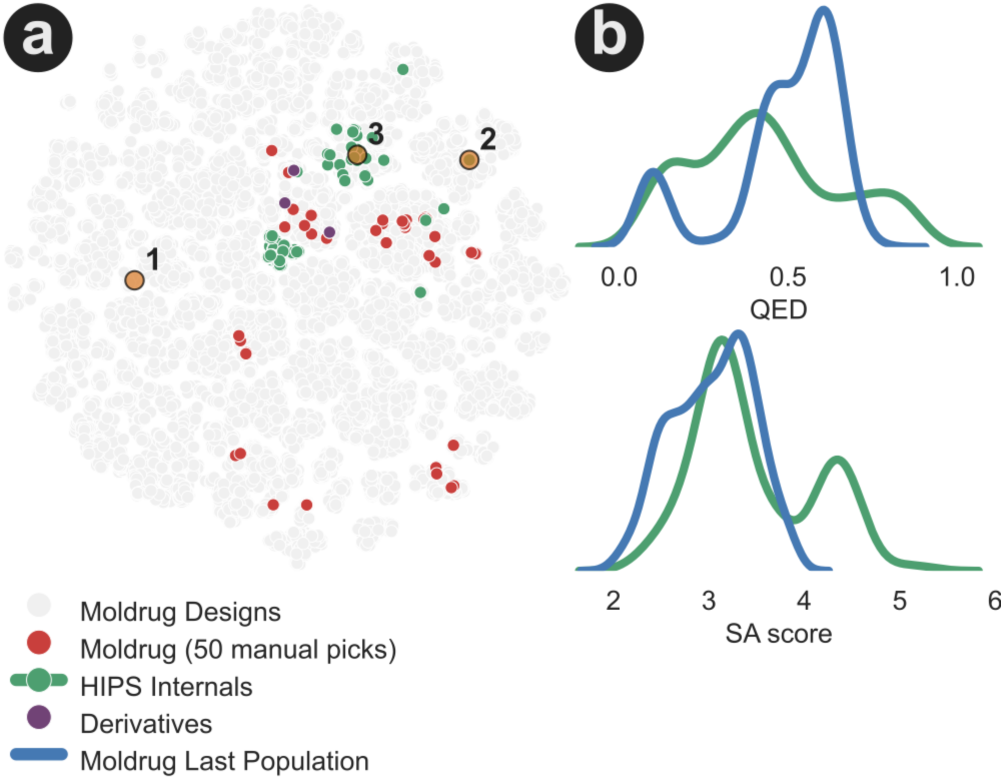
Coverage of chemical space. (a) t-distributed stochastic-neighbor embedding (t-SNE) analysis. The space was constructed based on the Morgan fingerprint of radius 2 and 2048 bits as implemented in RDKit^64^. After calculating the first 50 principal components (PCs), which explain 66.1% of the variance, the two main components of the t-SNE were derived from these PCs. Seed molecules (brown), HIPS internal molecules (green), Moldrug-designed molecules (gray), and the 50 manually selected molecules (red) are shown. (b) Distribution of QED and SA score for the internal HIPS dataset of LasB inhibitors (green) and the 870 molecules of the last populations of all campaigns (blue).

Moldrug employs an evolutionary algorithm to navigate chemical space, evolving a population of optimized candidates rather than converging on a single solution. The final populations across all campaigns yielded 870 unique molecules. To evaluate the quality of these generated molecules, we compared their Quantitative Estimate of Drug-likeness (QED) and Synthetic Accessibility (SA) scores against the internal HIPS dataset (Figure 3b).

The QED score ranges from 0 to 1, with higher values indicating superior drug-likeness. While the internal HIPS dataset covers the full QED spectrum, it is notably skewed toward lower values, likely due to the presence of highly polar phosphonate groups within many compounds. In contrast, the 870 molecules generated by Moldrug exhibited a distinct bimodal distribution. Most of the 870 molecules form a broad peak at intermediate-to-high QED values, with approximately 30% of the molecules achieving a QED > 0.6. This represents a significant optimization milestone as the initial seed molecules **1**, **2**, and **3** possess low QED values of 0.4, 0.4, and 0.2, respectively. This transition demonstrates Moldrug’s capacity to evolve scaffolds with potentially improved pharmacological profiles, effectively overcoming the inherent limitations of the low-QED starting point. A secondary, smaller peak at lower QED is primarily attributed to molecules derived from seed **3**. The inherent structural complexity of seed **3**, combined with the inclusion of the hydroxamic acid moiety, was naturally penalized by the QED score. Despite this, the presence of these molecules is vital, as the hydroxamic acid is functionally critical for target binding, suggesting that these results remain highly relevant for practical drug design despite the lower numerical score.

The SA score ranges from 1 (easy to synthesize) to 13 (highly difficult). The internal HIPS dataset displays a bimodal distribution with a minor peak at intermediate complexity. In comparison, the 870 molecules generated by Moldrug show a favorable shift toward lower values, maintaining a mean SA score of approximately 3. This shift indicates that the optimized candidates are not “virtual curiosities” but are predicted to be as accessible, if not more so, than the molecules already synthesized within the HIPS dataset. The highest SA scores (indicating greater complexity) are concentrated in campaigns with seed **3**. This correlates with the seed’s high initial complexity, suggesting that while Moldrug explores more intricate scaffolds when prompted, it successfully maintains a threshold of synthetic feasibility across the population.

From the 870 molecules generated by Moldrug, a chemically diverse subset of 50 molecules was manually selected to maximize both predicted performance and coverage of the explored chemical space (Figure 3a). These molecules were then subjected to alchemical absolute binding free energy calculations using BindFlow.

### Binding free energy ranking

Prior to prospective evaluation, the absolute binding free energy protocol was retrospectively validated on the three seed molecules for which both experimental IC_50_ values and protein–ligand complex structures were available^53,59,65^. The results, summarized in Table 1, show good agreement between calculated and experimental affinities.

**Table 1.**
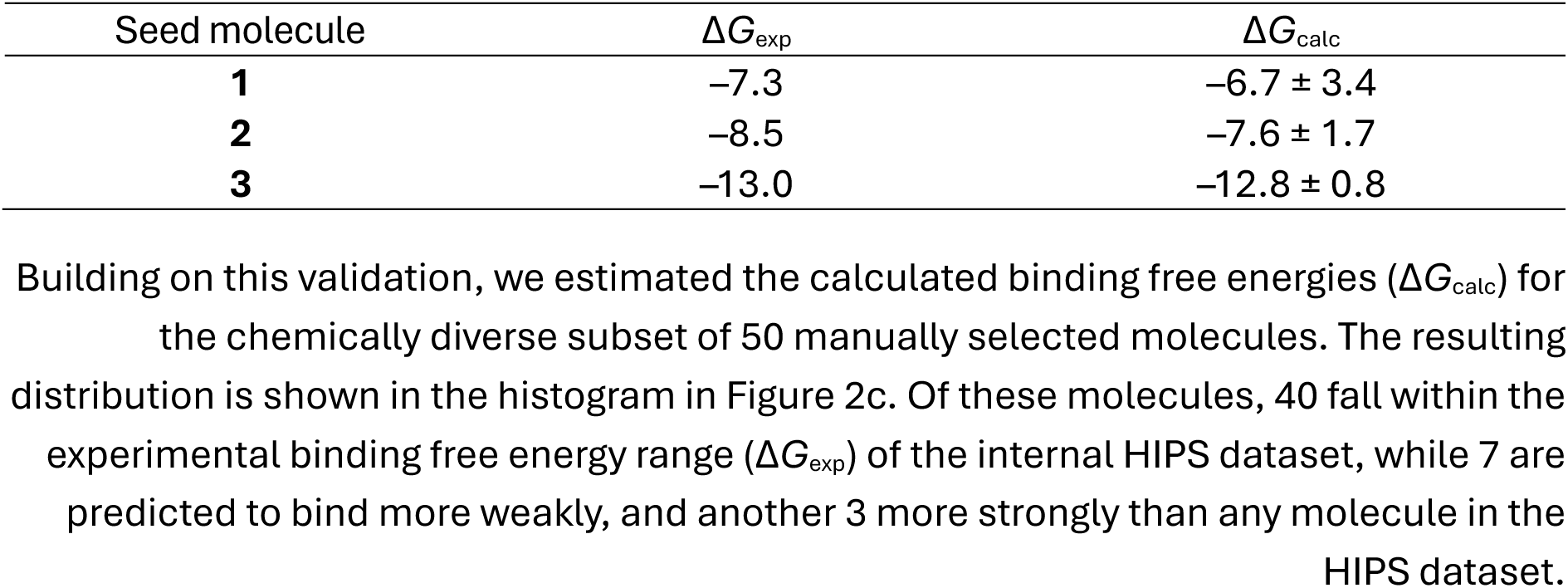
Retrospective study for the three seed molecules employed in this study. ΔG_exp_ and ΔG_calc_ are expressed in kcal/mol. ΔG_exp_ was obtained from the IC_50_ reported either in the original reference^53,57,59,65^ or from the internal HIPS dataset.

Notably, the uncertainty for seed **1** is substantial. Because this error corresponds to the standard error of the mean (SEM) across three independent replicates, it may indicate inherent instability within this protein–ligand complex during the simulations.

### Hit optimization

Molecules **2b** and **2c** (**Error! Reference source not found.**a) showed the highest predicted potency, however, they were not progressed to synthesis due to practical considerations regarding project scope and synthetic complexity. Nevertheless, a detailed analysis of these unique structures is provided in the Supporting Information to guide future investigations into these underexplored regions of the binding pocket.

The following two most potent predicted molecules, **3a** and **3b** (Figure 4), introduce novel non-*Zn*^2+^-binding scaffolds by chemically simplifying seed **3** molecule, thereby increasing ligand efficiency (the ratio of binding affinity to molecular mass). Consequently, these scaffolds were prioritized for further hit optimization.

**Figure 4.**
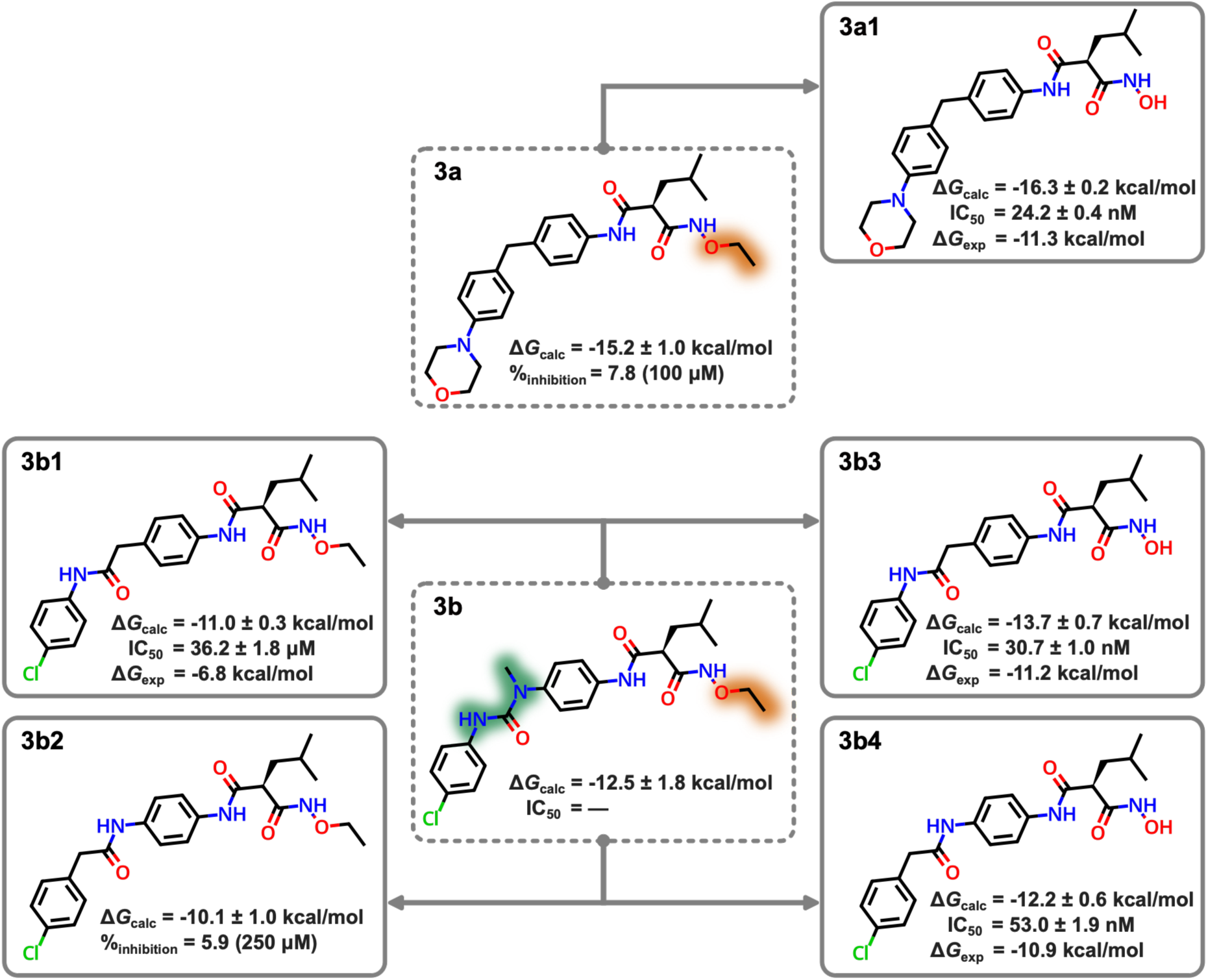
Structure–activity relationship (SAR) data. Moldrug-designed molecules are shown in dotted boxes, while derivative compounds are shown in solid boxes. The calculated binding free energy (ΔG_calc_) is reported alongside the experimental IC_50_ values when available, as well as the corresponding derived experimental binding free energy (ΔG_exp_). For compounds **3a** and **3b2** the percent of inhibition of LasB at the specific compound concentration is reported.

### Hit optimization strategy

With hit **3b** molecule identified, we sought to determine if the central urea linker could be replaced without disrupting target binding. We explored the binding tolerance toward corresponding amide analogues as isosteric replacements. Both hit molecules, **3a** and **3b**, introduced an interesting *O*-alkylation of the hydroxamic acid, yielding an *O*-ethyl hydroxamate. If tolerated, this modification could be exploited to occupy the remaining volume of the binding pocket.

However, given the critical role of the hydroxamic acid in LasB inhibition and in view of a potential metabolic liability of the *O*-ethyl hydroxamate, the corresponding derivatives without the *O*-alkylation (*i.e.*, free hydroxamic acids) were also subjected to absolute binding free energy calculations using BindFlow. In total, five new derivatives were computationally evaluated (Figure 4).

Comparison of the Δ*G*_calc_ for the pairs **3a1**/**3a**, **3b3**/**3b1**, and **3b4**/**3b2**; shows that *O*-ethyl alkylation consistently reduces binding affinity, resulting in losses of 1.1, 2.7, and 2.1 kcal/mol, respectively. This decrease can be rationalized by several factors: (i) the removal of the hydroxyl hydrogen, which participates in a key hydrogen bond with Glu141 (see Figure 2b); (ii) a decrease in the negative charge on the hydroxamic oxygen atom (–0.43 for the hydroxyl group versus –0.27 for the ethoxy group in **3b1** and **3b2**, and –0.22 in **3a**), which weakens electrostatic interactions with the *Zn*²⁺ ion; and (iii) alterations in solvation effects, as suggested by visual inspection of the simulation trajectories. Despite their weaker predicted binding relative to the corresponding hydroxamic acids, the *O*-ethyl hydroxamates retained robust calculated affinities, providing a compelling rationale for biological evaluation.

The isosteric replacement of the urea group in the **3b** series (comparing **3b** against **3b1** and **3b2**) unfavorably impacts binding; notably, this reduction is only tolerated if the O-alkylation of the hydroxamic acid is absent. Additionally, comparison of the pairs **3b1**/**3b2** and **3b3**/**3b4** (Figure 4) indicates that the relative position of the amide nitrogen in the **3b** series contributes approximately 1 kcal/mol to the binding affinity. Collectively, these computational results motivated the synthesis of both the *O*-ethyl hydroxamates and their free hydroxamic acid counterparts.

### Synthesis of the novel compounds

All chlorophenyl compounds were synthesized via a common synthetic route using different starting materials (Scheme 1). Initial amide coupling reactions were performed using T3P as the coupling agent under basic conditions, affording the desired amides (**p1** and **p5**) after aqueous workup without the need for further purification. Subsequent Boc deprotection under acidic conditions yielded the corresponding free amines (**p2** and **p6**), which were isolated following basic workup. Introduction of the isobutyl moiety, which was previously shown to be critical for biological activity, was achieved under the same amide-coupling conditions, affording the corresponding malonate ester intermediates (**p3** and **p7**). Conversion of the esters to the corresponding free hydroxamic acids (**3b3** and **3b4**) was accomplished by treatment with hydroxylamine in the presence of catalytic potassium cyanide over extended reaction times. For the synthesis of the structurally extended derivatives (**3b1** and **3b2**), ester saponification afforded the corresponding carboxylic acids, which subsequently underwent amide coupling to yield the *N*-ethoxy-2-isobutylmalonamides.

**Scheme 1.**
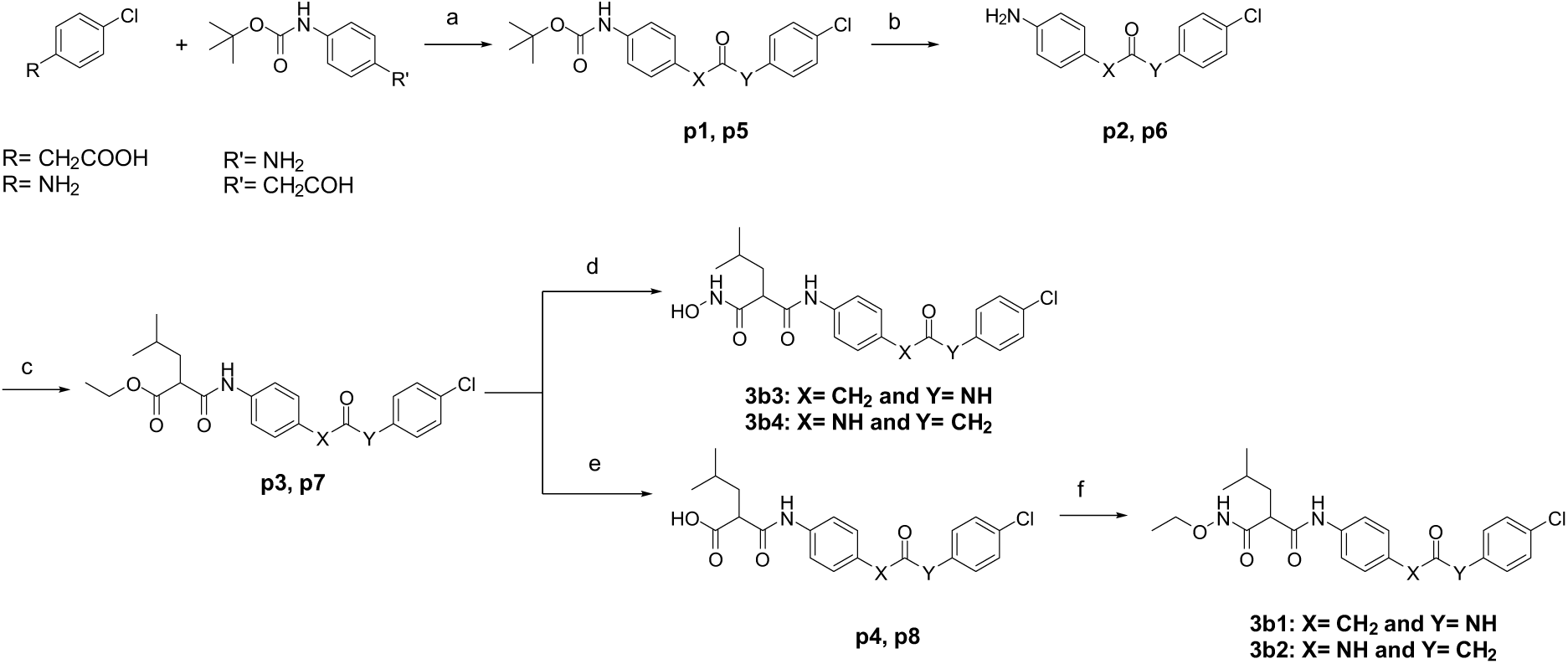
Synthesis of the four chlorophenyl compounds (3b3, 3b4, 3b1 and 3b2) incorporating the free and extended hydroxamic acid moieties. Reagents and conditions: (a) Et_3_N, T3P (50% in EtOAc), CH_2_Cl_2_, 0 °C to r.t., 16 h, 52–95%, (b) TFA, CH_2_Cl_2_, r.t., 20–60 min, 89–100% (c) Et_3_N, T3P (50% in EtOAc), CH_2_Cl_2_, 0 °C to r.t., 16 h, 89–100% (d) NH_2_OH, KCN, 1:1 THF/MeOH, r.t., 18–96 h, 32–100%, (e) 1 M LiOH in 1:1 THF/ H_2_O, r.t., 2–18 h, 56–62% (f) Et_3_N, T3P (50% in EtOAc), CH_2_Cl_2_, 0 °C to r.t., 16 h, 90–96%.

The synthetic route to the morpholinobenzyl compounds **3a** and **3a1** commenced with a Suzuki–Miyaura cross-coupling reaction employing an XPhos ligand, which facilitated efficient coupling while minimizing the formation of by-products (Scheme 2). This was followed by acidic Boc deprotection and subsequent amide coupling under basic conditions using T3P as the coupling reagent affording compound **p12**. Formation of the key ester intermediate **p12** enabled divergence into two synthetic pathways. Conversion to the free hydroxamic acid **3a1** was achieved by treatment of the ester with hydroxylamine in the presence of catalytic potassium cyanide over extended reaction time. In the second pathway, ester saponification yielded the corresponding carboxylic acid **p13**, which subsequently underwent amide coupling to give the structurally extended *N*-ethoxy-2-isobutylmalonamide derivative **3a**.

**Scheme 2.**
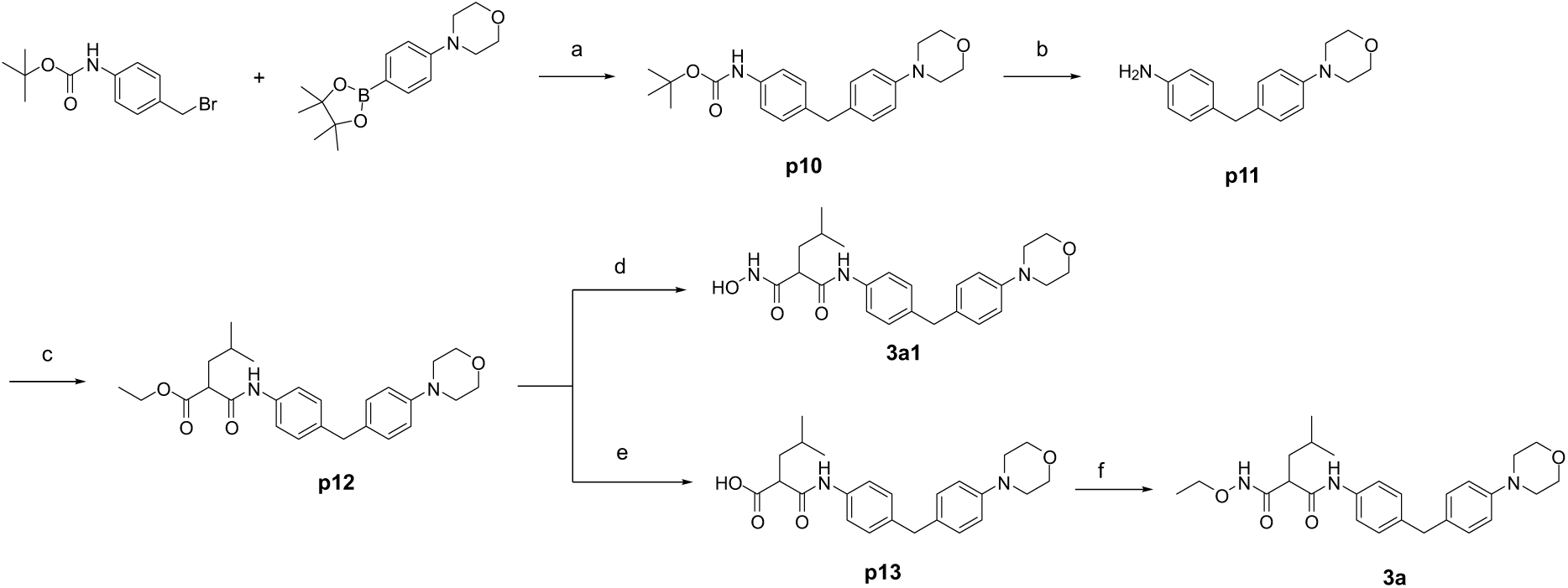
Synthesis of the two morpholinobenzyl compounds (3a1 and 3a) with the free and structurally extended hydroxamic acid moiety. Reagents and conditions: (a) [Pd_2_(dba)_3_], XPhos, K_3_PO_4_, 1:1 1,4-dioxane/ H_2_0, 100 °C, 1 h, MW, 50%. (b) TFA, CH_2_Cl_2_, r.t., 90 min, 40% (c) Et_3_N, T3P (50% in EtOAc), CH_2_Cl_2_, 0 °C to r.t., 16 h, 81% (d) NH_2_OH, KCN, 1:1 THF/MeOH, r.t., 72 h, 68%, (e) 1 M LiOH in 1:1 THF/ H_2_O, r.t., 90 min, 100% (f) Et_3_N, T3P (50% in EtOAc), CH_2_Cl_2_, 0 °C to r.t., 16 h, 90%.

### *In vitro* evaluation and structure–activity relationships (SAR)

The *in vitro* inhibitory activity of the newly synthesized compounds was evaluated using the functional FRET-based assay, as previously established^36^. The natural product phosphoramidon was employed as a reference inhibitor. IC_50_ values and percentage of inhibition results were determined from at least two independent experiments performed in duplicate. IC_50_ values were converted to binding free energies (Δ*G*_exp_) as described above and compared with ΔG_calc_.

#### Computational *vs* experimental binding affinity

Although the computational model overestimated absolute binding affinities by approximately 1 kcal/mol to 5 kcal/mol, the relative ranking of the tested molecules demonstrated significant consistency (Figure 4), yielding a Kendall rank correlation coefficient of 0.6.

In structure-based drug design, accurate rank-ordering is often more critical than absolute precision, as it directly informs lead prioritization. Consequently, these results remain highly valuable for guided decision-making. Notably, this systematic overestimation was not observed for the initial seed molecules (Table 1), suggesting a potential shift in predictive behavior within the newly explored chemical space.

#### In vitro results

The experimental evaluation confirmed the qualitative trends predicted computationally for both *O*-ethyl alkylation and amide-nitrogen repositioning. Among the six compounds tested, the structurally extended hydroxamic acid derivatives **3b2**, **3b1**, and **3a** exhibited little to no inhibitory activity, in agreement with prior studies^53^, highlighting the critical role of hydrogen-bonding interactions with Glu141. Compounds **3b2** and **3a** showed less than 10% inhibition at a concentration of 100 μM, while compound **3b1** displayed an IC₅₀ value of 36 μM. This potency was markedly reduced relative to the corresponding free hydroxamic acid derivatives. Consequently, these three compounds were not progressed to further ADMET (absorption, distribution, metabolism, excretion, and toxicity) evaluation.

In contrast, the free hydroxamic acid derivatives demonstrated excellent inhibitory activity. The morpholine derivative **3a1** exhibited an IC₅₀ of 24.2 ± 0.4 nM, while the chlorinated compounds **3b3** and **3b4** showed IC₅₀ values of 30.7 ± 1.0 nM and 53.0 ± 1.9 nM, respectively. These values are comparable to previously reported inhibitors^59^, while the lower molecular weights of these compounds contribute to improved ligand efficiency.

#### *In vitro* ADMET profiling

Given our goal of identifying compounds that have favorable drug-like properties, we determined the *in vitro* ADMET profile of the three nanomolar binders (compounds **3a1**, **3b3** and **3b4**). All tested compounds exhibited excellent metabolic and plasma stability in both murine and human fractions (Table 2). While the seed molecules, specially **2** and **3**, showed minimal cytotoxicity at 100 µM (**Error! Reference source not found.**), the generated compounds displayed moderate cytotoxicity against both HepG2 and A549 cells. Yet, this still gives a good selectivity window in view of the potent on-target activity. Chromatographic logD_7.4_ and plasma protein binding (PPB) were similar for the three compounds. While the *in vitro* profile of the three compounds is generally comparable, there is a minor advantage of **3b4** with slightly higher unbound fraction and slightly reduced cytotoxicity.

**Table 2.**
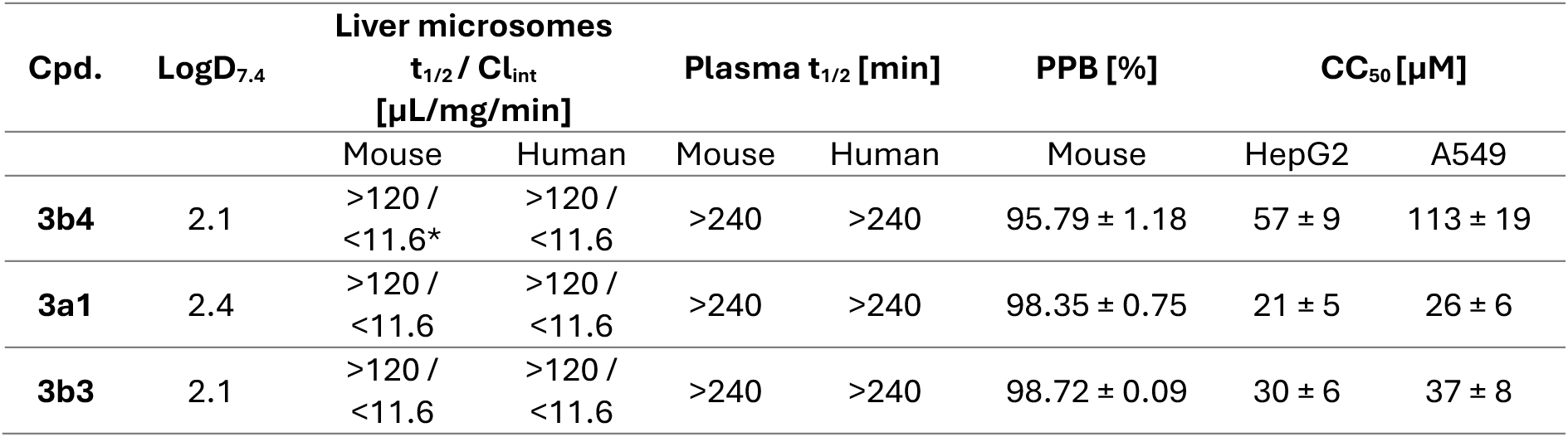
In vitro ADMET profile of 3b4, 3a1, and 3b3. Values represent mean ± standard deviation of at least two independent replicates. LogD7.4 was determined using a chromatographic method. t_1/2_: half-life, Cl_int_: intrinsic clearance, PPB: plasma protein binding; CC_50_: concentration leading to 50% cell viability. *66 ± 8% remaining after 2 h.

| Cpd. | $\log D_{7.4}$ | Liver microsomes | | Plasma $t_{1/2}$ [min] | | PPB [%] | $CC_{50}$ [μM] | |
| --- | --- | --- | --- | --- | --- | --- | --- | --- |
| | | $t_{1/2} / Cl_{int}$<br>[μL/mg/min] | | | | | | |
|  |  | Mouse | Human | Mouse | Human | Mouse | HepG2 | A549 |
| **3b4** | 2.1 | >120 /<br><11.6* | >120 /<br><11.6 | >240 | >240 | 95.79 ± 1.18 | 57 ± 9 | 113 ± 19 |
| **3a1** | 2.4 | >120 /<br><11.6 | >120 /<br><11.6 | >240 | >240 | 98.35 ± 0.75 | 21 ± 5 | 26 ± 6 |
| **3b3** | 2.1 | >120 /<br><11.6 | >120 /<br><11.6 | >240 | >240 | 98.72 ± 0.09 | 30 ± 6 | 37 ± 8 |

## Discussions and conclusions

In this study, we demonstrate a strategy for the prospective discovery of bioactive small molecules by integrating fragment-based *de novo* design with alchemical ABFE calculations. Applied to the virulence-associated metalloprotease LasB from *P. aeruginosa*, the workflow enabled the prioritization of candidates from a diverse, computationally generated chemical space, leading to the identification of three novel nanomolar inhibitors with favorable metabolic profiles. Beyond the specific application to LasB, these findings show that the coupling of generative molecular design with quantitative affinity prediction focuses experimental efforts on high-value regions of chemical space. Since Moldrug and Bindflow are distributed under free licenses and with extensive documentation, the workflow may be readily adopted by the community to alternative targets.

The Moldrug framework offers distinct advantages by accommodating a wide range of drug-relevant objectives to guide the optimization process^29^. While this study employed the Derringer–Suich desirability functions^66^ to aggregate QED^61^, the SA score^62^ and the Vina docking score^60^, the framework is explicitly extensible. Properties relevant to Gram-negative drug design, such as outer membrane permeability descriptors, efflux pump evasion proxies^67^ or ADMET profiles^68,69^, may be readily incorporated into Moldrug as additional desirability components without architectural changes to the optimizer. This modularity provides a practical advantage over approaches with fixed objectives, allowing the campaign to be steered towards design constraints that most relevant to the target indication. Furthermore, the degree of molecular novelty is tunable: tight scaffold constraints approximate lead optimization, whereas unconstrained exploration yields genuinely novel chemotypes.

In contrast to traditional virtual screening, which relies on databases of pre-existing compounds, the Moldrug algorithm generates novel molecules with potentially advantageous intellectual property profiles. To regulate structural complexity, we employed the SA score, which is computationally efficient, well-validated for drug-like molecules in known chemical space^62^, and crucial for managing molecular generation within Moldrug. However, several candidates in this study received acceptable SA scores yet presented substantial synthetic challenges. Consequently, human expertise remains essential as a final post-hoc filter to select candidates for subsequent development. Integrating retrosynthesis-based alternatives, such as SCScore^70^, AiZynthFinder^71^ or the recently proposed FSscore^72^ represent straightforward extensions that could reduce this gap in future iterations of the pipeline.

The workflow presented here is economically efficient. The costs for synthesis of a novel small molecule may range from hundreds to several thousands of USD depending on structural complexity and the synthetic route. Furthermore, biophysical binding measurements, such as isothermal titration calorimetry or surface plasmon resonance, involve additional costs in instrument time, protein consumption, and compound quantity. Given current accessible cloud computing prices, the computational cost of running BindFlow ABFE calculations is approximately 100 USD or less per compound, representing a fraction of the cost of a failed synthesis–assay cycle. If even a modest fraction of weak binders is computationally filtered prior to experimental validation, the approach is highly cost-effective.

Our findings reveal that bulky hydrophobic groups (compounds **2b** and **2c**) may be tolerated in the Zn^2+^-binding region, offering a path for ligand elaboration into unexplored chemical space. To ensure the reliability of these future studies, it will be essential to employ advanced force field descriptions of metal–ligand environments to minimize non-systematic errors in binding predictions.

Considering that ranking by ABFE let to nanomolar binders, this study demonstrates that ABFE is suitable for prioritizing not on congeneric but also diverse chemical sets. However, despite the theoretical rigor of ABFE, its practical accuracy is sensitive to the proposed binding pose^20,73^. For the *de novo* molecules generated here, poses were obtained via constrained docking. Because MD simulations within the ABFE protocol are too short to spontaneously correct a potentially inaccurate pose, uncertainties from the docking protocol propagate into the computed free energies, likely explaining the reduced ABFE accuracy observed for the novel compounds compared to the seed molecules. This constitutes, in our view, the most consequential practical limitation of the current pipeline and, generally, of prospective ABFE studies on diverse compound sets. Thus, in scenarios involving high chemical diversity and binding pose uncertainty, ABFE results are best interpreted as relative rankings within chemical clusters rather than as absolute binding free energies. Improving the binding pose generation will be essential to bridge the gap between computational prediction and experimental validation.

In conclusion, our results underscore the unique capability of computational *de novo* design to explore chemical hypotheses that are often impractical to pursue via traditional experimental routes. While the synthesis of unconventional or high-risk scaffolds is frequently cost-prohibitive, Moldrug enables rapid, cost-effective interrogation of these architectures, identifying novel directions for drug development. Rather than offering a single candidate, this approach provides a rational framework for navigating promising regions of chemical space. When integrated with the high-precision ranking of BindFlow and experimental validation, it establishes a powerful, complementary strategy for efficient and informed drug discovery. Using this pipeline, we identified three novel nanomolar LasB inhibitors with excellent metabolic stability; future optimization may focus on mitigating cytotoxicity.

## Methods

### Structure retrieving and processing

Three different protein-ligand complexes served as starting point for the drug design campaign: PDB ID: **6FZX**^53^, PDB ID: **GGM4**^58^, and **32KK**^57^. Their corresponding ligands served as seeds for Moldrug^29^ and will be denoted on this research as **1**, **2**, and **3**, respectively for simplicity.

All protein–ligand complexes were aligned in PyMOL^74^ for easy post-processing. Crystal waters, calcium ions and ligands were removed, and the protein was protonated at pH = 7 with the PDB2PQR software^75^. The PDBQT file of the protein with the *Zn*^2+^ ion was generated by the *prepare_receptor* script from ADFRsuite-1.0^76^ followed by the *zinc_pseudo.py*^77^ script to detect and configurate tetrahedral *Zn* atoms. However, *Zn* tetrahedral atoms were not detected for 6FZX. Docking grids were constructed with a spacing of 0.375 Å; center at (23.5, 62, 27.5) Å and 66 (X), 53 (Y), and 66 (Z) grid points. For 6FZX, the center was placed at (21.5, 62, 27.5) Å and 56 grid points in the X direction were used instead of 66. Affinity maps were generated with the script *prepare_gpf4zn.py*^77^ and the AutoDock4Zn^78^ force fields. AutoDock Vina 1.2.0^60,77^ was used as docking engine alongside with Moldrug^29^. The protein PDB file with the *Zn*^2+^ and *Ca*^2+^ ions was generated and used as input for the MD simulations.

### Design strategies and parameters for fragment-based optimization

*Zn*-metalloproteins represent challenging targets for docking algorithms, as *Zn*^2+^–ligand interactions are often poorly modeled. In addition to employing the improved AutoDock4Zn^78^ force fields, we maintained a constant *Zn*^2+^-binding moiety for the majority of Moldrug campaigns. This approach assumes that any systematic error in modeling *Zn*^2+^–ligand interactions would affect all designed molecules equally, thus preserving the reliability of relative ranking.

Fragment-based optimization was performed using Moldrug (v3.7.2)^29^ in seven independent campaigns. Each campaign was conducted in triplicate and divided into three or four stages. Three campaigns were initiated from seed molecules **1** and **2**, and one campaign from molecule **3**. The population size was set to 50 molecules with a selection pressure of 1 and a child proportion of 0.5. The conformation of generated molecules was constrained to their respective seed structure using the *local_only* Moldrug mode, with 100 conformers and an exhaustiveness of 9. The Cost fitness function, which integrates AutoDock-Vina (Vina) score^60^, reflecting molecular potency, quantitative estimation of drug-likeness (QED)^61^, and synthetic accessibility (SA) score^62^. These properties were combined into four different sets of desirability functions (see Table 3).

**Table 3.** Definition of Derringer–Suich desirability functions^66^ used in the seven campaigns. Here l, t and u denote the lower, target and upper values, respectively, while r and w are exponents controlling the weighting of each contribution—Vina score, QED and SA score—to the weighted geometric mean (D).

| Identification | Vina score |  |  |  | QED |  |  |  | SA score |  |  |  |
| --- | --- | --- | --- | --- | --- | --- | --- | --- | --- | --- | --- | --- |
| | $t$ | $u$ | $r$ | $w$ | $l$ | $t$ | $r$ | $w$ | $t$ | $u$ | $r$ | $w$ |
| D-1 | -14.0 | -2.0 | 1.0 | 1.0 | 0.4 | 0.8 | 0.5 | 1.0 | 3.0 | 7.0 | 0.5 | 1.0 |
| D-2 | -14.0 | -4.0 | 1.0 | 1.0 | 0.4 | 0.8 | 1.0 | 1.0 | 2.0 | 7.0 | 1.0 | 1.0 |
| D-3 | -14.0 | -2.0 | 1.0 | 1.0 | 0.4 | 0.8 | 1.0 | 1.0 | 2.0 | 7.0 | 1.0 | 1.0 |
| D-4 | -14.0 | -2.0 | 1.0 | 1.0 | 0.5 | 1.0 | 0.5 | 1.0 | 3.0 | 7.0 | 0.5 | 1.0 |

Moldrug relies on CReM^8,79^ as the molecular generator. We employed the published CReM database of interchangeable fragments with maximum synthetic accessibility score of 2.^8,79^ Table 4 summarizes the two protocols employed in this study. The four-stage protocol (Protocol 1) follows the approach described in Ref. ^29^, inspired by the progression from lead generation to lead optimization. Protocol 1 was applied to all campaigns initiated from seeds **1** and **2**. Protocol 2 omits the first stage (01_grow)—as **3** does not allow further structure expansion—and employs 45 generations on the 02_allow_grow stage (corresponding to 01-02_allow_grow on Protocol 2).

**Table 4.** Protocol definitions inspired by Ref^23^. CReM parameters and number of generations at each stage.

| Protocol | Stage | Radius | min_size | max_size | min_inc | max_inc | Generations |
| --- | --- | --- | --- | --- | --- | --- | --- |
| P-1 | 01_grow | 3 | 0 | 0 | 1 | 6 | 20 |
|  | 02_allow_grow | 3 | 0 | 2 | -2 | 4 | 25 |
|  | 03_pure_mutate | 3 | 1 | 8 | -5 | 3 | 40 |
|  | 04_local | 3 | 0 | 1 | -1 | 1 | 15 |
| P-2 | 01-02_allow_grow | 3 | 0 | 2 | -2 | 4 | 45 |
|  | 03_pure_mutate | 3 | 1 | 8 | -5 | 3 | 40 |
|  | 04_local | 3 | 0 | 1 | -1 | 1 | 15 |

Table 5 summarizes the seven campaigns. The combination of different seeds, desirability functions, and CReM parameters, along with three replicates per campaign, was designed to explore a diverse chemical space. In all campaigns, heavy atoms involved in *Zn*^2+^ binding were protected to reduce potential errors in describing *Zn*^2+^-ligand interactions, except for campaign V, which permits modification of the hydroxyl groups in the hydroxamic acid. Campaigns III and VII additionally protect the hydrogen atom of the NH–OH group.

**Table 5.**
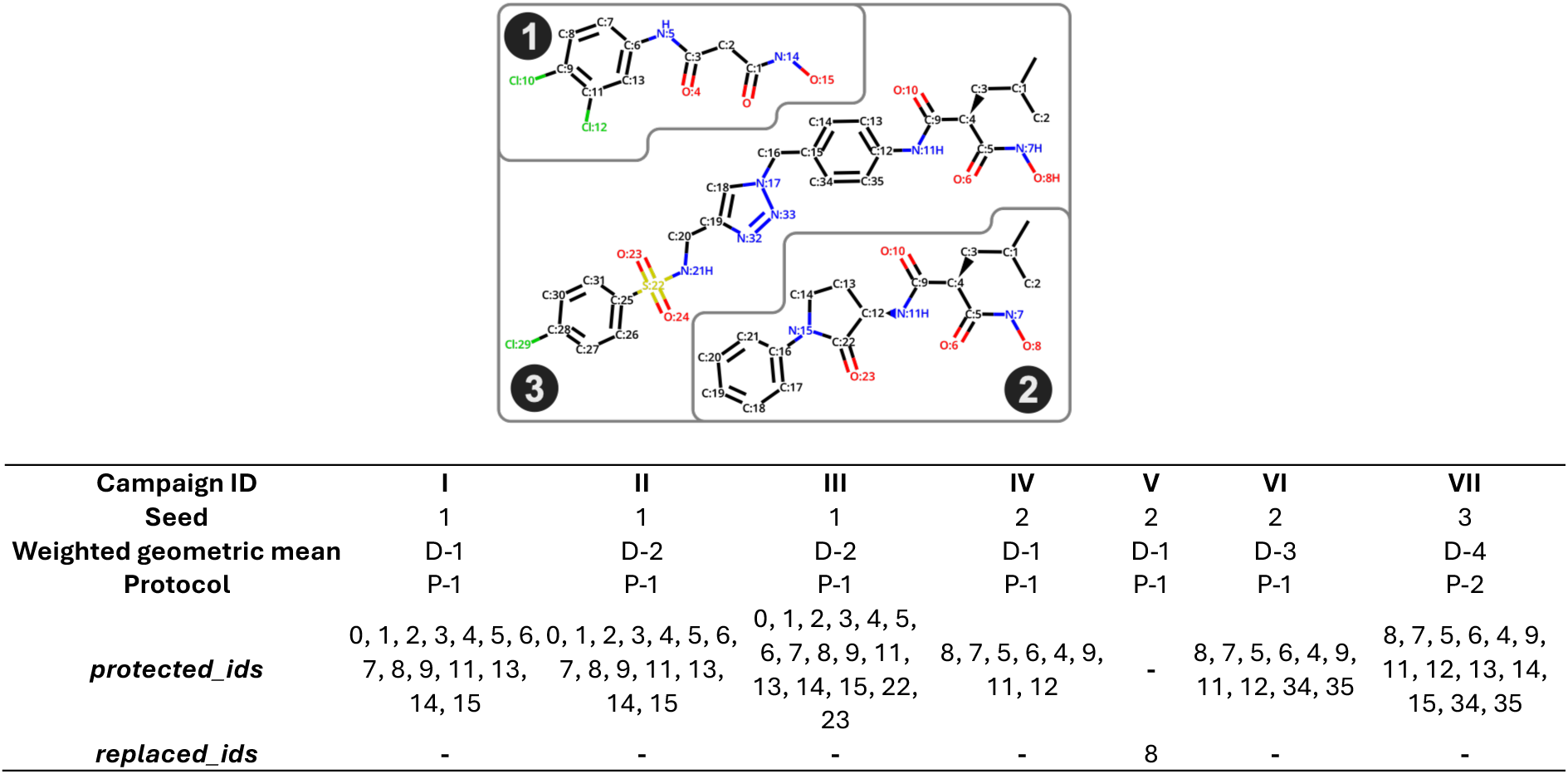
Definition of each campaign. The table summarizes the protocol used (see Table 4) and the applied weighted geometric mean (see Table 3), and additional CReM parameters specific for each campaign. It also presents atoms either protected (protected_ids) or submitted (replace_ids) for structural modification to CReM as well the employed seed molecule. Atom indices for each seed molecule are provided for reference.

### Binding affinity prediction and prioritization

The seven triplicated campaigns yielded a total of 1050 molecules in the last generation. After removing duplicates, the 870 remaining molecules were visually inspected. A final selection of 50 molecules was made to ensure representation from each seed molecule. Specifically, 8, 33 and 9 molecules originating from the seed molecules **1**, **2** and **3**, respectively, were selected. Selection was based on the desirability scores, chemical representativeness, and practical synthetic criteria. These 50 molecules were subjected to ABFE calculation at FEP level of theory.

Inhibition constants (*K_i_*) were calculated from experimentally determined *IC*_50_ values using the Cheng–Prusoff equation^80^:

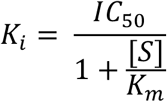

where [*S*] is the substrate concentration and *K_m_* is the Michaelis constant of the substrate for LasB. The substrate concentration used in the assay was 150 μM, while the *K_m_* value for LasB was taken from Nishino and Powers^36^ and reported as 110 μM.

Experimental binding free energies, which are defined for the reaction

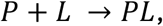

where *P* is the protein, *L* the ligand and *PL* the protein-ligand complex, were estimated according to

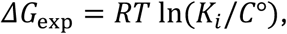

where *R* is the universal gas constant, *C*° is the standard concentration of 1 M, and *T* is the absolute temperature equals to 310.15 K for all calculations.

#### Free energy perturbation calculations

Alchemical absolute binding free energies (FEP) calculations were performed with BindFlow-0.13.0.post24^12^. The default BindFlow protocol for soluble proteins was applied. In brief, a total of 43 lambda windows for the complex and 22 lambda windows for the ligand in water were used, where each window was simulated for over 10 ns during production. Non-default or additional relevant settings are described below. Amber14sb^81^ force field topologies for the protein with the *Zn*^2+^ and *Ca*^2+^ cofactors were generated via OpenMM^82^ and ParmEd^83^. Ligands interactions were described with OpenFF-2.0.0^84^, and their topologies were generated within BindFlow using the TOFF Python library^85^. The TIP3P water model^86^ and AMBER ion parameters^87^ were employed. Hydrogen mass repartitioning^88,89^ was used with a factor of 2.5. Three independent replicates were performed for each ligand if not stated otherwise. The reported Δ*G*_calc_ values and associated errors correspond to the mean across replicates and the standard error of the mean. The Multistate Bennett Acceptance Ratio (MBAR)^90^ method was used as the free energy estimator. GROMACS-2022.4^91^ was used as molecular-dynamics engine.

## Data availability statement

Moldrug and BindFlow are free software hosted at https://github.com/ale94mleon/moldrug and https://github.com/ale94mleon/BindFlow respectively. Scripts and input files required for reproducing our results and analysis are accessible at: https://doi.org/10.5281/zenodo.23234680.

## Acknowledgments

This research was funded by the European Union. Views and opinions expressed are however those of the author(s) only and do not necessarily reflect those of the European Union. Neither the European Union nor the granting authority can be held responsible for them. This project has received funding from the European Union’s Horizon Europe Research and Innovation Program under the Marie Skłodowska-Curie Grant Agreement No. 860592 and No. 101081463 and by the Deutsche Forschungsgemeinschaft (DFG, German Research Foundation) via grant INST 256/539-1.

