## Supporting Information PDF for "Prospective discovery of LasB inhibitors by integrating generative design, free energy ranking, and experimental validation"

#### Supporting results

Molecules **2b** and **2c** showed the highest predicted potency from the Moldrug-designed molecules. These two molecules introduce structural modifications to the $Zn^{2+}$-binding scaffold that extend into a previously underexplored region of the binding pocket, which appears to accommodate bulkier hydrophobic substituents (Figure S1a).

Because the initial three replicates displayed relatively large statistical uncertainties (SEM), and because binding free energies lower than –15 kcal/mol may indicate pathological behavior in alchemical binding free energy calculations[1], we increased the sampling to ten replicates for these ligands to obtain more reliable free-energy estimates.

We hypothesized that the enhanced binding of these scaffolds arises from two synergistic effects: (i) direct $Zn^{2+}$–$N$ coordination introduced by the nitrogen atom in the bulky substituent, and (ii) partial dehydration of the $Zn^{2+}$ coordination sphere promoted by steric shielding of the metal center. In particular, we reasoned that bulkier hydrophobic groups extending toward the solvent-exposed region could displace coordinated water molecules, thereby favoring stronger $Zn^{2+}$–ligand interactions.

To test this hypothesis, we analyzed the solvation state of the $Zn^{2+}$ center using cumulative radial distribution functions (RDFs), $Zn^{2+}$–$O_{water}$. The plateau observed in the cumulative RDFs (Figure S1b) indicates the presence of one or two coordinating water molecules within ~0.23 nm in certain molecules and simulation replicates. We further quantified the first hydration shell using a 0.23 nm cutoff (Figure S1b). Accordingly, a cumulative RDF value of zero at 0.23 nm corresponds to a dehydrated $Zn^{2+}$ center, whereas nonzero values indicate the presence of coordinated water molecules.

To disentangle the effects of steric bulk and nitrogen coordination, we designed a set of control compounds. First, we included **2a**, which places a hydrophobic substituent in the same region of the binding pocket as **2b** and **2c** but with reduced steric bulk, leaving the $Zn^{2+}$ center more solvent-exposed. Second, for each of the three parent scaffolds (**2a**, **2b**, and **2c**), we generated derivatives in which the nitrogen atom in the substituent was either removed or introduced. This design allowed us to independently assess the contributions of steric shielding and *Zn–N* coordination.

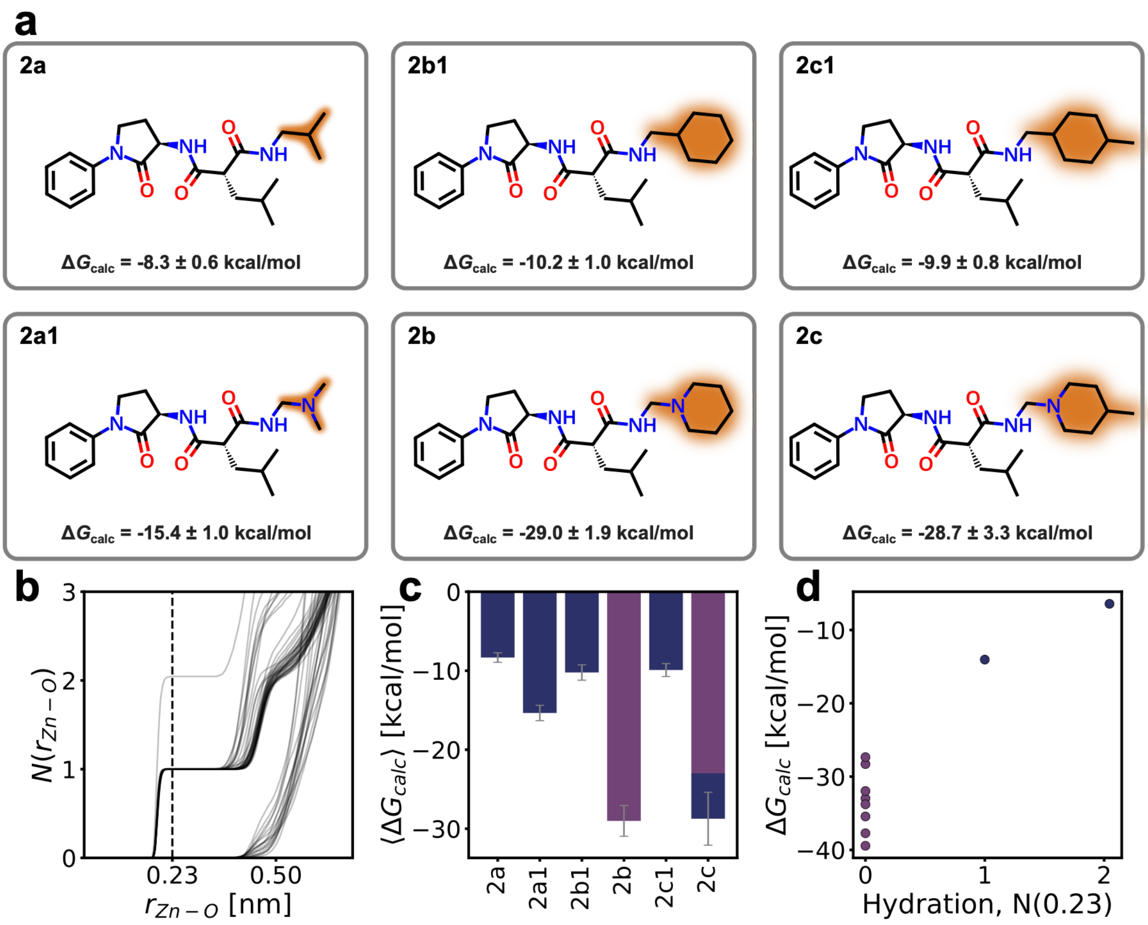


*Figure S1.* ***Hydration state of the Zn$^{2+}$ coordination sphere and its relationship to binding free energy*****.** ***(a)*** *Chemical structures of the investigated ligands.* ***(b)*** *Cumulative radial distribution functions, N($r_{Zn-O}$), computed from classical MD simulations for all ligands and replicates, the last 10 ns of equilibration were used. The dashed line indicates the cutoff (0.23 nm) used to define the hydration first-shell.* ***(c)*** *Mean calculated binding free energies ($\langle \Delta G_{calc} \rangle$) for the six selected ligands (ten replicates were used). Bars are colored according to the fraction of replicates classified as dry (purple) or hydrated (blue) based on the hydration criterion in panel (a). Error bars represent the standard error of the mean across replicates.* ***(d)*** *Replica-resolved binding free energies for ligand* ***2c*** *plotted as a function of hydration number at the cutoff distance. Purple points correspond to dry replicates and blue points to hydrated replicates.*

For the less bulky **2a** scaffold, the *Zn*$^{2+}$ center remains largely solvent-exposed and retains approximately one coordinated water molecule across all replicates, irrespective of nitrogen substitution (Figure S1c). In this series, incorporation of a nitrogen atom improves binding free energy without significantly altering *Zn*$^{2+}$ hydration, indicating that the affinity gain primarily originates from strengthened *Zn*$^{2+}$*–N* coordination rather than desolvation of the *Zn*$^{2+}$.

In contrast, for the bulkier **2b** and **2c** scaffolds, nitrogen substitution is associated with displacement of the coordinated water molecule, resulting in a predominantly dehydrated *Zn*$^{2+}$ center and substantially stronger predicted binding (Figure S1c). Thus, in these systems, affinity enhancement appears to arise from a combined effect of direct metal coordination and favorable *Zn*$^{2+}$ desolvation. Notably, **2c** displays replicadependent hydration behavior: replicates that retain a coordinated water molecule systematically exhibit weaker binding free energies than those in which the $Zn^{2+}$ center is dehydrated (Figure S1d), directly linking incomplete desolvation to reduced affinity.

Together, these results support a mechanistic model in which nitrogen incorporation strengthens binding through direct $Zn^{2+}$ coordination across all scaffolds, while steric bulk modulates the hydration state of the metal center. Only for sufficiently bulky scaffolds does dehydration provide an additional energetic contribution to binding.

Despite their promising predicted binding profiles, these compounds were not prioritized for synthesis due to practical considerations related to synthetic feasibility and project scope. Nevertheless, their behavior highlights structurally underexplored regions of the binding pocket that may merit future investigation. Further studies employing more advanced models for $Zn^{2+}$ coordination could help refine the characterization of these scaffolds.

*Table S1. **Cytotoxicity of seed compounds1, 2 and 3 in HepG2 cells at 100 µM**. Cell viability was assessed after exposure to each compound at a fixed concentration (100 µM). **1** showed moderate cytotoxicity, reducing HepG2 viability to 60 ± 6%, whereas **2** and **3** exhibited minimal cytotoxicity, with cell viability remaining above 90% under the same conditions.*

| Compound | HepG2 viability at 100 µM [%] |
|---|---|
| **1** | 60 ± 6 |
| **2** | >90[2] |
| **3** | >90 |

### Synthesis

#### Chemicals, materials and methods

NMR experiments were performed on a Bruker Ultrashield plus 500 (500 MHz) spectrometer. Spectra were acquired at 300 K, using deuterated dimethylsulfoxide (DMSO-$d_6$) or deuterated chloroform ($CDCl_3$) as solvents. Chemical shifts for $^1H$ and $^{13}C$ spectra were recorded in parts per million (ppm) using the residual non-deuterated solvent as the internal standard (for DMSO-$d_6$: 2.50 ppm, $^1H$; 39.52 ppm, $^{13}C$; for $CDCl_3$: 7.27 ppm, $^1H$; 77.00, $^{13}C$). Coupling constants ($J$) are given in Hertz (Hz). Data are reported as follows: chemical shift, multiplicity (s = singlet, d = doublet, t = triplet, m = multiplet, br = broad and combinations of the aforementioned) coupling constants and integration.

Mass spectrometry was performed on a SpectraSystems-MSQ LCMS system (Thermo Fisher, Dreieich, Germany) using a Hypersil Gold column, 100 × 2.1 mm, 1.9 µm. At a flow rate of 600 µL/min, a gradient of $H_2O$ (0.1% FA) and ACN (0.1% FA) was used, starting from 5% ACN and then increased to 100% over 5 min. The mass spectrum was measured in positive and negative ionization modes in a range from 100–600 *m/z*. The UV spectrum was recorded at 254 nm. An EC HPLC Column (Analytical), NUCLEODUR C18 Pyramid, 3 µm, 150 × 2 mm was used for separation. At a flow rate of 250 µL/min, the gradient of (A)

$H_2O$ + 0.1% FA and (B) ACN + 0.1% FA was held at 10% B for 1 min and then increased to 100% B over 4 min. It was held there for 1.5 min before the gradient was decreased to 10% B over 0.3 min where it was held for 0.7 min, resulting in a total method time of 7.5 min. The mass spectrum was measured in positive mode in a range from 120–1000 m/z. UV detection was performed at 254 nm.

Compounds were purified by either Preparative high-performance liquid chromatography (HPLC) or flash column chromatography.

Flash chromatography was performed using the automated flash chromatography system CombiFlash NextGen 300+ (Teledyne Isco, Lincoln, NE, USA) equipped with RediSepRf silica columns (Axel Semrau, Sprockhövel Germany). Preparative HPLC was performed with a ThermoScientific Ultimate 3000 system using the column NUCLEODUR C18 Gravity, 250 × 16 mm, 5 µm Macherey-Nagel (Düren Germany). At a flow rate of 10 mL/min, the gradient of $H_2O$ (0.05% FA) and ACN (0.05% FA) starting from 5% ACN and then increased to 100% over 40 min. The UV spectrum was recorded at 254 nm. All final compounds showed purity of > 95% according to LC-MS data.

#### General Procedures

##### *General Procedure A: Amide Coupling*

A solution of the carboxylic acid (1.0 equiv) in $CH_2Cl_2$ (0.1–0.2 M) containing $Et_3N$ (3.0 equiv) was cooled to 0 °C in an ice bath. Propanephosphonic acid anhydride (T3P, 2.0–3.0 equiv, 50 wt % solution in EtOAc)) was added dropwise, followed by addition of the amine (1.2–2.0 equiv). The reaction mixture was allowed to warm to room temperature and stirred for 16 h. Upon completion of the reaction, the mixture was diluted with EtOAc and washed sequentially with $H_2O$, saturated aqueous $NaHCO_3$, and brine. The combined organic layers were dried over $MgSO_4$, filtered, and concentrated under reduced pressure. The crude product was either used directly in the subsequent step without further purification or purified by flash column chromatography to afford the desired product.

##### *General Procedure B: Boc Deprotection*

The Boc-protected intermediate (1.0 equiv) was dissolved in dry $CH_2Cl_2$ and the solution was cooled in an ice bath. Trifluoroacetic acid (TFA) was added dropwise, and the reaction mixture was allowed to warm to room temperature and stirred for up to 1 h. Upon completion, the reaction mixture was quenched with aqueous $NaHCO_3$ and extracted with EtOAc. The combined organic layers were washed with brine, dried over $MgSO_4$, filtered, and concentrated under reduced pressure. The crude product was purified by flash column chromatography to afford the desired product.

##### *General Procedure C: Hydroxaminolysis*

The ester derivatives were dissolved in a 1:1 mixture of tetrahydrofuran and methanol (THF/MeOH). Aqueous hydroxylamine solution (50%, 11.0–33.0 equiv) and a catalytic amount of potassium cyanide (0.2 equiv) were added sequentially, and the reaction mixture was stirred at room temperature for 2–7 d. Upon completion, the solvents were removed under reduced pressure, and the residue was diluted with $H_2O$. The product was extracted with EtOAc, and the combined organic layers were dried over $MgSO_4$, filtered, and concentrated under reduced pressure. The crude product was purified by preparative HPLC or flash column chromatography to afford the desired product.

##### *General Procedure D: Saponification*

A solution of the ester in THF (0.225 M) was treated with aqueous LiOH (2 M, 0.225 M) and the reaction mixture was stirred at room temperature for 2 h. The solvents were removed under reduced pressure, and the residue was acidified to pH 2 with 1 M HCl. The product was extracted with EtOAc, and the combined organic layers were washed with $H_2O$ and brine, dried over $MgSO_4$, filtered, and concentrated under reduced pressure. The crude product was used in the subsequent step without further purification.

tert-Butyl (4-(2-(4-chlorophenyl)acetamido)phenyl)carbamate (**p1**)

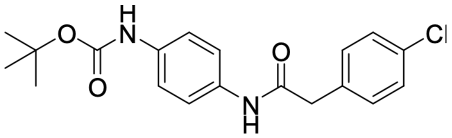


Compound **p1** was synthesized according to General Procedure A, using 2-(4-chlorophenyl)acetic acid (330 mg, 1.93 mmol), tert-butyl (4-aminophenyl)carbamate (443 mg, 2.13 mmol), T3P (50 % in EtOAc, 2.28 mL, 3.87 mmol), and $Et_3N$ (808 µL, 5.80 mmol) in $CH_2Cl_2$ (9.67 mL) for 16 h. Compound **p1** was obtained as a crude product (360 mg, 52%) and used in the subsequent step without further purification. LC $t_R$ = 4.61 min, MS (ESI) *m/z:* 361.22 $[M+H]^+$. $^1$H NMR (500 MHz, *DMSO-$d_6$*) δ 10.04 (s, 1 H), 9.24 (br s, 1 H), 7.42 - 7.47 (m, 2 H), 7.36 (s, 6 H), 3.60 (s, 2 H), 1.46 (s, 9 H). $^{13}$C NMR (126 MHz, DMSO-$d_6$) δ 168.24, 152.78, 135.09, 135.01, 133.57, 131.21, 130.99 (2C), 128.18 (2C), 119.61 (2C), 118.40 (2C), 78.83, 42.33, 28.13 (3C).

*N*-(4-aminophenyl)-2-(4-chlorophenyl)acetamide (**p2**)

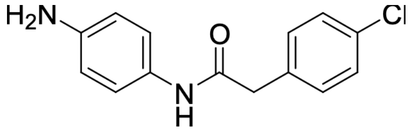


Compound **p2** was synthesized according to General Procedure B, using carbamate **p1** (360 mg, 0.998 mmol), TFA (1.7 mL, 22.2 mmol) in dry $CH_2Cl_2$ (6.65 mL) at room temperature for 20 min. Compound **p2** was obtained as a crude product (231 mg, 89 %) and was used in the next step without further purification. LC $t_R$= 2.73 min, MS (ESI) *m/z:* 261.13 $[M+H]^+$. $^1$H NMR (500 MHz, DMSO-$d_6$) δ 9.74 (s, 1H), 7.41 – 7.29 (m, 4H), 7.20 (d, *J* = 8.5 Hz, 2H), 6.48 (d, *J* = 8.5 Hz, 2H), 4.86 (br s, 2H), 3.55 (s, 2H). $^{13}$C NMR (126 MHz,

DMSO-$d_6$) δ 168.09, 145.31, 135.92, 131.59, 131.39 (2C), 128.73, 128.63 (2C), 121.36 (2C), 114.24 (2C), 42.79.

Ethyl 2-((4-(2-(4-chlorophenyl)acetamido)phenyl)carbamoyl)-4-methylpentanoate (**p3**)

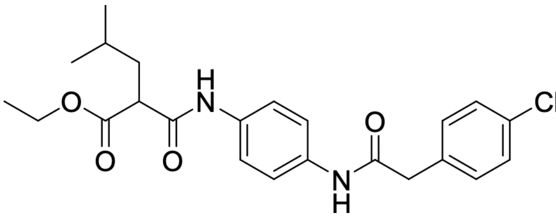

Compound **p3** was synthesized according to General Procedure A, using amine **p2** (75 mg, 0.287 mmol), 2-(ethoxycarbonyl)-4-methylpentanoic acid (108 mg, 0.575 mmol), T3P (50% in EtOAc, 0.34 mL, 0.575 mmol), and $Et_3N$ (0.12 mL, 0.862 mmol) in $CH_2Cl_2$ (1.44 mL) for 16 h. Compound **p3** was obtained as a crude product (107 mg, 86%) and used in the subsequent step without further purification. LC $t_R$= 4.67 min, MS (ESI) *m/z:* 431.25 $[M +H]^+$. $^1$H NMR (500 MHz, DMSO-$d_6$) δ 10.15 (br d, *J* = 9.8 Hz, 2H), 7.56 – 7.45 (m, 4H), 7.44 – 7.30 (m, 4H), 4.16 – 4.04 (m, 2H), 3.62 (s, 2H), 3.57 – 3.51 (m, 1H), 1.78 – 1.70 (m, 1H), 1.69 – 1.61 (m, 1H), 1.53 – 1.45 (m, 1H), 1.16 (t, *J* = 7.3 Hz, 3H), 0.93 – 0.86 (m, 6H) $^{13}$C NMR (126 MHz, DMSO-$d_6$) δ 170.29, 168.89, 167.19, 135.51, 135.34, 134.77, 131.72, 131.49 (2C), 128.68 (2C), 120.18 (2C), 120.02 (2C), 61.07, 51.33, 42.81, 37.92, 26.12, 23.13, 22.43, 14.51.

*N*1-(4-(2-(4-chlorophenyl)acetamido)phenyl)-*N*3-hydroxy-2-isobutylmalonamide (**3b4**)

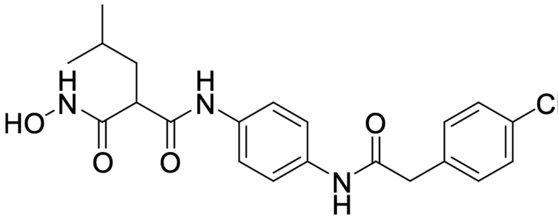

Compound **3b4** was synthesized according to General Procedure C using compound **p3** (25 mg, 0.058 mmol), hydroxylamine (50% in water, 78 µL, 1.28 mmol), potassium cyanide (0.75 mg, 0.011 mmol) in a 1:1 mixture of THF/MeOH (0.29 mL) for 96 h. The crude product was purified by flash column chromatography on silica gel affording compound **3b4** (24 mg, 100 %). LC $t_R$= 3.75 min, MS (ESI) *m/z:* 418.19 $[M +H]^+$. $^1$H NMR (500 MHz, DMSO-$d_6$) δ 10.54 (br s, 1H), 10.13 (s, 1H), 9.71 (s, 1H), 8.96 (br s, 1H), 7.50 (s, 4H), 7.39 (s, 4H), 3.62 (s, 2H), 3.19 (br t, *J* = 7.5 Hz, 1H), 1.67 (br t, *J* = 7.2 Hz, 2H), 1.51 – 1.44 (m, 1H), 0.87 (br dd, *J* = 6.4, 3.5 Hz, 6H) $^{13}$C NMR (126 MHz, DMSO-$d_6$) δ 168.88, 167.92, 166.83, 166.83, 135.51, 135.20, 134.76, 131.72, 131.49, 128.68, 120.22, 119.92, 50.28, 42.83, 38.50, 26.16, 22.91, 22.72. $^{13}$C NMR (126 MHz, DMSO-$d_6$) δ 168.88, 167.92, 166.83, 135.51, 135.20, 134.76, 131.72, 131.49 (2C), 128.68 (2C), 120.22 (2C), 119.92 (2C), 50.28, 42.83, 38.50, 26.16, 22.91, 22.72.

2-((4-(2-(4-Chlorophenyl)acetamido)phenyl)carbamoyl)-4-methylpentanoic acid (**p4**)

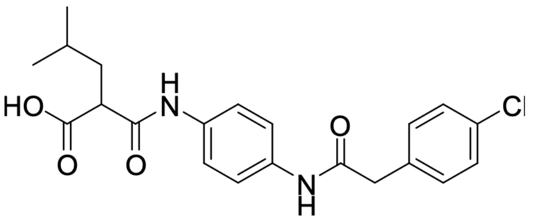

Compound **p4** was synthesized according to General Procedure D using ester **p3** (220 mg, 0.51 mmol) in THF (2.2 mL) and aqueous LiOH (2 M, 2.2 mL). The reaction mixture was stirred at room temperature for 2 h and compound **p4** was obtained without further purification (128 mg, 62%). LC $t_R$ = 4.14 min, MS (ESI) *m/z:* 403.19 $[M +H]^+$. $^1$H NMR (500 MHz, DMSO-$d_6$) δ 12.59 (br s, 1H), 10.16 – 10.06 (m, 2H), 7.57 – 7.46 (m, 4H), 7.43 – 7.30 (m, 4H), 3.62 (br d, *J* = 4.1 Hz, 2H), 3.27 – 3.22 (m, 1H), 1.76 – 1.68 (m, 1H), 1.56 – 1.43 (m, 2H), 0.89 – 0.86 (m, 6H). $^{13}$C NMR (DMSO-$d_6$, 126 MHz) δ : 171.9, 171.5, 168.9, 167.7, 135.5, 135.2, 134.9, 131.7, 131.5, 128.7, 120.0, 120.0, 51.4, 50.4, 42.8, 37.8, 37.7, 26.2, 26.1, 23.2, 22.6, 22.4.

*N*1-(4-(2-(4-chlorophenyl)acetamido)phenyl)-*N*3-ethoxy-2-isobutylmalonamide (**3b2**)

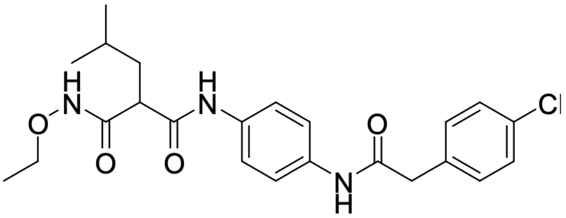

Compound **3b2** was synthesized according to General Procedure A using carboxylic acid **p4** (47 mg, 0.116 mmol), *O*-ethylhydroxylamine hydrochloride (13.66 mg, 0.140 mmol), T3P (50% in EtOAc, 206 µL, 0.349 mmol), and $Et_3N$ (48 µL, 0.349 mmol) in $CH_2Cl_2$ (0.58 mL). The reaction mixture was stirred for 16 h. The crude product was purified by flash column chromatography on silica gel to afford compound **3b2** as a beige powder (50 mg, 96%). LC $t_R$ = 4.11 min, MS (ESI) *m/z:* 446.27 $[M + H]^+$. $^1$H NMR (500 MHz, DMSO-$d_6$) δ 10.96 (s, 1H), 10.12 (s, 1H), 9.72 (s, 1H), 7.54 – 7.46 (m, 4H), 7.41 – 7.32 (m, 4H), 3.79 (br d, *J* = 7.0 Hz, 2H), 3.65 – 3.59 (m, 2H), 3.19 (t, *J* = 7.6 Hz, 1H), 1.68 (t, *J* = 7.0 Hz, 2H), 1.48 (dt, *J* = 13.3, 6.6 Hz, 1H), 1.12 (t, *J* = 7.0 Hz, 3H), 0.92 – 0.84 (m, 6H) $^{13}$C NMR (126 MHz, DMSO-$d_6$) δ 168.42, 167.23, 166.25, 135.04, 134.79, 134.26, 131.24, 131.03 (2C), 128.22 (2C), 119.86 (2C), 119.48 (2C), 70.55, 50.00, 42.35, 37.88, 25.79, 22.37, 22.24, 13.37.

tert-Butyl (4-(2-((4-chlorophenyl)amino)-2-oxoethyl)phenyl)carbamate (**p5**)

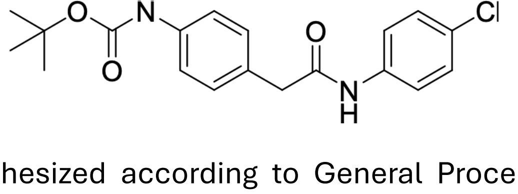

Compound **p5** was synthesized according to General Procedure A using 2-(4-((tert-butoxycarbonyl)amino)phenyl)acetic acid (400 mg, 1.59 mmol), 4-chloroaniline (223 mg,

1.75 mmol), T3P (50% in EtOAc, 1.88 mL, 3.18 mmol), and $Et_3N$ (665 µL, 4.78 mmol) in $CH_2Cl_2$ (7.96 mL). The reaction mixture was stirred for 2 h. Compound **p5** (550 mg, 95%) was obtained and used in the next step without further purification. LC $t_R$ = 5.25 min, MS (ESI) *m/z:* 361.22 [M +H]$^+$. $^1$H NMR (DMSO-$d_6$, 500 MHz) δ 10.23 (s, 1H), 9.28 (br s, 1H), 7.6-7.6 (m, 2H), 7.38 (br d, 2H, *J*=8.4 Hz), 7.3-7.4 (m, 2H), 7.2-7.2 (m, 2H, *J*=8.5 Hz), 3.54 (s, 2H), 1.46 (s, 9H). $^{13}$C NMR (126 MHz, DMSO-$d_6$) δ 169.50, 152.80, 138.20, 138.09, 129.29 (2C), 129.26, 128.62 (2C), 126.70, 120.63 (2C), 118.15 (2C), 78.93, 42.62, 28.14 (3C).

2-(4-Aminophenyl)-N-(4-chlorophenyl)acetamide (**p6**)

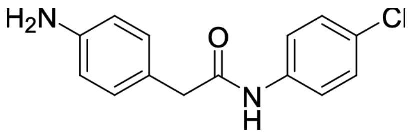


Compound **p6** was synthesized according to General Procedure B using carbamate **p5** (550 mg, 1.52 mmol) and TFA (3 mL, 39.2 mmol) in $CH_2Cl_2$ (15 mL). The reaction mixture was stirred at room temperature for 1 h. Compound **p6** (397 mg, quantitative) was obtained and used in the next step without further purification. LC $t_R$ = 3.00 min, MS (ESI) *m/z:* 261.17 [M +H]$^+$. $^1$H NMR (DMSO-$d_6$, 500 MHz) δ 10.14 (s, 1H), 7.6-7.6 (m, 2H), 7.3-7.4 (m, 2H), 6.96 (d, 2H, *J*=8.2 Hz), 6.5-6.5 (m, 2H), 4.93 (s, 2H), 3.41 (s, 2H). $^{13}$C NMR (DMSO-$d_6$, 126 MHz) δ 170.6, 147.7, 138.8, 129.9, 129.0, 127.0, 123.0, 121.0, 114.3, 43.1.

Ethyl 2-((4-(2-((4-chlorophenyl)amino)-2-oxoethyl)phenyl)carbamoyl)-4-methylpentanoate (**p7**)

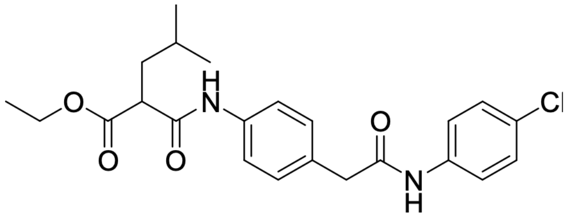


Compound **p7** was synthesized according to General Procedure A using amine **p6** (397 mg, 1.52 mmol), 2-(ethoxycarbonyl)-4-methylpentanoic acid (573 mg, 3.05 mmol), T3P (50% in EtOAc, 1.79 mL, 3.05 mmol), and $Et_3N$ (636 µL, 4.57 mmol) in $CH_2Cl_2$ (15.2 mL). The reaction mixture was stirred for 16 h. Compound **p7** was obtained as a crude product (656 mg, quantitative) and used in the next step without further purification. LC $t_R$ = 5.35 min, MS (ESI) *m/z:* 431.33 [M +H]$^+$. $^1$H NMR (500 MHz, DMSO-$d_6$) δ 10.29 – 10.24 (m, 1H), 10.21 (s, 1H), 7.64 – 7.59 (m, 2H), 7.52 (d, *J* = 8.5 Hz, 2H), 7.36 – 7.32 (m, 2H), 7.26 (d, *J* = 8.5 Hz, 2H), 4.14 – 4.05 (m, 2H), 3.58 (s, 2H), 3.57 – 3.53 (m, 1H), 1.79 – 1.71 (m, 1H), 1.69 – 1.60 (m, 1H), 1.54 – 1.44 (m, 1H), 1.15 (t, *J* = 7.2 Hz, 3H), 0.88 (dd, *J* = 8.8, 6.6 Hz, 6H). $^{13}$C NMR (126 MHz, DMSO-$d_6$) δ 170.28, 169.83, 167.44, 138.64, 137.88, 131.41, 129.92 (2C), 129.09 (2C), 127.19, 121.11 (2C), 119.76 (2C), 61.09, 51.38, 43.15, 37.88, 26.12, 23.13, 22.42, 14.50.

*N*1-(4-(2-((4-chlorophenyl)amino)-2-oxoethyl)phenyl)-*N*3-hydroxy-2-isobutylmalonamide (**3b3**)

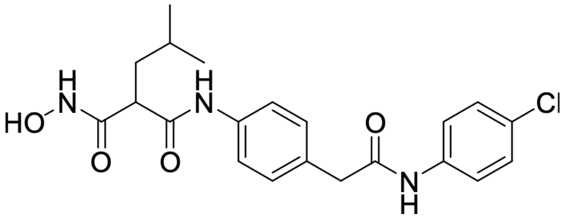

Compound **3b3** was synthesized according to General Procedure C using compound **p7** (300 mg, 0.69 mmol), hydroxylamine (50% in water, 1.41 mL, 23.0 mmol), potassium cyanide (9 mg, 0.14 mmol), in a 1:1 mixture of THF/MeOH (3.48 mL). The reaction mixture was stirred at room temperature for 18 h. The crude product was purified by flash column chromatography on silica gel to afford compound **3b3** as a white solid (92 mg, 32%). LC $t_R$ = 4.30 min, MS (ESI) *m/z:* 418.22 $[M +H]^+$. $^1$H NMR (500 MHz, DMSO-$d_6$) δ 10.55 (s, 1H), 10.26 (s, 1H), 9.74 (s, 1H), 8.97 (s, 1H), 7.61 (d, *J* = 8.9 Hz, 2H), 7.51 (d, *J* = 8.5 Hz, 2H), 7.37 – 7.32 (m, 2H), 7.24 (d, *J* = 8.4 Hz, 2H), 3.58 (s, 2H), 3.21 (t, *J* = 7.6 Hz, 1H), 1.71 – 1.63 (m, 2H), 1.53 – 1.42 (m, 1H), 0.87 (t, *J* = 6.7 Hz, 6H). $^{13}$C NMR (126 MHz, DMSO-$d_6$) δ 169.83, 168.13, 166.83, 138.63, 137.84, 131.24, 129.83 (2C), 129.09 (2C), 127.19, 121.11 (2C), 119.86 (2C), 50.36, 43.13, 38.54, 26.18, 22.88, 22.72.

2-((4-(2-((4-Chlorophenyl)amino)-2-oxoethyl)phenyl)carbamoyl)-4-methylpentanoic acid (**p8**)

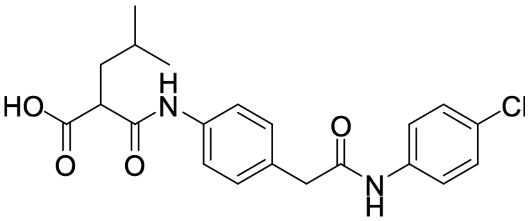

Compound **p8** was synthesized according to General Procedure D using ester **p7** (300 mg, 0.69 mmol) in THF (3 mL) and aqueous LiOH (2 M, 3 mL). The reaction mixture was stirred for 18 h. Compound **p8** was obtained without further purification (159 mg, 56%). LC $t_R$ = 4.72 min, MS (ESI) *m/z:* 403.22 $[M +H]^+$. $^1$H NMR (500 MHz, DMSO-$d_6$) δ 12.54 (br s, 1H), 10.28 (s, 1H), 10.16 (s, 1H), 7.64 – 7.60 (m, 2H), 7.54 (d, *J* = 8.4 Hz, 2H), 7.37 – 7.32 (m, 2H), 7.25 (d, *J* = 8.5 Hz, 2H), 3.58 (s, 2H), 3.47 (dd, *J* = 9.0, 5.8 Hz, 1H), 1.76 – 1.69 (m, 1H), 1.65 – 1.58 (m, 1H), 1.55 – 1.44 (m, 1H), 0.88 (dd, *J* = 8.9, 6.6 Hz, 6H). $^{13}$C NMR (126 MHz, DMSO-$d_6$) δ 171.90, 169.84, 168.03, 138.65, 138.05, 131.24, 129.88 (2C), 129.09 (2C), 127.18, 121.11 (2C), 119.65 (2C), 51.51, 43.15, 37.88, 26.15, 23.21, 22.42.

*N*1-(4-(2-((4-chlorophenyl)amino)-2-oxoethyl)phenyl)-*N*3-ethoxy-2-isobutylmalonamide (**3b1**)

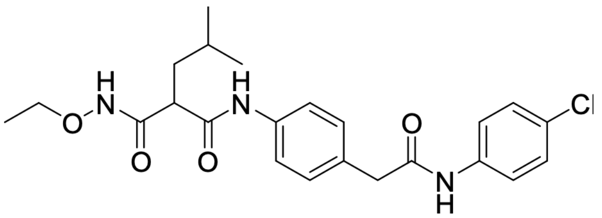

Compound **3b1** was synthesized according to General Procedure A using carboxylic acid **p8** (160 mg, 0.397 mmol), *O*-ethylhydroxylamine hydrochloride (46 mg, 0.48 mmol), T3P (50% in EtOAc, 351 µL, 1.19 mmol), and $Et_3N$ (166 µL, 1.19 mmol) in $CH_2Cl_2$ (1.99 mL). The reaction mixture was stirred for 16 h. The crude product was purified by flash column chromatography on silica gel to afford compound **3b1** as a white solid (158 mg, 90%). LC $t_R$ = 4.92 min, MS (ESI) *m/z:* 446.31 $[M + H]^+$. $^1H$ NMR (500 MHz, DMSO-$d_6$) δ 10.26 (s, 1H), 9.99 (s, 1H), 7.61 (d, *J* = 1.0 Hz, 2H), 7.52 (d, *J* = 8.5 Hz, 2H), 7.37 – 7.32 (m, 2H), 7.24 (d, *J* = 8.5 Hz, 2H), 3.76 (br t, *J* = 6.9 Hz, 1H), 3.63 (s, 2H), 3.58 (s, 2H), 3.08 (s, 3H), 1.76 – 1.69 (m, 1H), 1.63 – 1.51 (m, 2H), 1.23 (br s, 1H), 0.89 (dd, *J* = 10.5, 6.4 Hz, 6H). $^{13}C$ NMR (126 MHz, DMSO-$d_6$) δ 169.85, 168.64, 138.64, 138.10, 131.15, 129.83 (2C), 129.09 (2C), 127.18, 121.10 (2C), 119.86 (2C), 61.64, 48.90, 43.15, 38.50, 26.20, 22.99, 22.94, 14.56.

tert-Butyl (4-(4-morpholinobenzyl)phenyl)carbamate (**p11**)

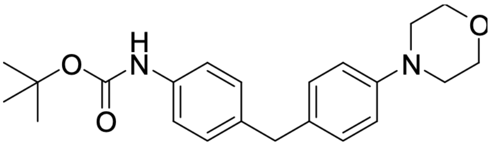

A microwave vial was charged with tert-butyl (4-(bromomethyl)phenyl)carbamate (300 mg, 1.04 mmol) and 4-(4-(4,4,5,5-tetramethyl-1,3,2-dioxaborolan-2-yl)phenyl)morpholine (356 mg, 1.24 mmol, 1.20 equiv) in a 9:1 mixture of 1,4-dioxane/$H_2O$ (6.10 mL). The mixture was degassed with argon, followed by the addition of $K_3PO_4$ (440 mg, 2.07 mmol, 2.00 equiv), $Pd_2(dba)_3$ (57 mg, 0.062 mmol, 6 mol %), and XPhos (49 mg, 0.103 mmol, 10 mol %). The reaction mixture was heated at 100 °C under microwave irradiation for 1 h. After completion of the reaction, the solvent was removed under reduced pressure, and the product was extracted with EtOAc. The combined organic layers were washed sequentially with $H_2O$ and brine, dried over $MgSO_4$, filtered, and concentrated under reduced pressure. The crude product was purified by flash column chromatography to afford compound **p11** (194 mg, 50%). LC $t_R$ = 4.89 min, MS (ESI) *m/z:* 369.34 $[M +H]^+$. $^1H$ NMR (500 MHz, DMSO-$d_6$) δ 9.21 (br s, 1H), 7.44 – 7.41 (m, 1H), 7.34 – 7.30 (m, 2H), 7.07 – 7.02 (m, 3H), 6.86 – 6.82 (m, 2H), 3.74 (s, 2H), 3.73 – 3.69 (m, 4H), 3.04 – 3.00 (m, 4H), 1.45 (s, 9H). $^{13}C$ NMR (DMSO-$d_6$, 126 MHz) δ 153.3, 149.8, 137.8, 135.9, 135.0, 132.8, 129.5, 129.4, 129.2, 128.9, 128.7, 128.6, 128.0, 127.2, 118.7, 115.8, 79.3, 66.6, 55.4, 49.2, 49.1, 28.6.

4-(4-Morpholinobenzyl)aniline (**p12**)

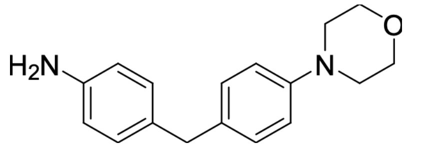

Compound **p12** was synthesized according to General Procedure B using carbamate **p11** (263 mg, 0.713 mmol) and TFA (1.75 mL, 22.9 mmol) in $CH_2Cl_2$ (7 mL). The reaction mixture was stirred at room temperature for 1.5 h. to yield compound **p12** (78 mg, 40%) as the crude product and was used in the next step without further purification. LC $t_R$ = 2.46 min, MS (ESI) *m/z:* 269.26 $[M+H]^+$. $^1H$ NMR (500 MHz, DMSO-$d_6$) δ 7.02 (d, *J* = 8.5 Hz, 2H), 6.83 (dd, *J* = 8.5, 1.9 Hz, 4H), 6.46 (d, *J* = 8.2 Hz, 2H), 4.83 (br s, 2H), 3.76 – 3.66 (m, 4H), 3.64 (s, 2H), 3.05 – 2.99 (m, 4H). $^{13}C$ NMR (126 MHz, DMSO-$d_6$) δ 149.64, 146.98, 133.67, 129.46 (2C), 129.46 (2C), 129.31, 115.75 (2C), 114.43 (2C), 66.61 (2C), 49.31 (2C), 40.04.

Ethyl 4-methyl-2-((4-(4-morpholinobenzyl)phenyl)carbamoyl)pentanoate (**p13**)

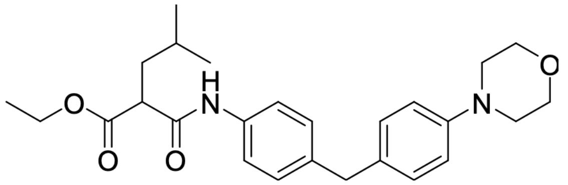

Compound **p13** was synthesized according to General Procedure A using amine **p12** (75 mg, 0.279 mmol), 2-(ethoxycarbonyl)-4-methylpentanoic acid (105 mg, 0.558 mmol), T3P (50% in EtOAc, 0.329 mL, 0.559 mmol), and $Et_3N$ (116 µL, 0.838 mmol) in $CH_2Cl_2$ (2.79 mL). The reaction mixture was stirred for 16 h. Compound **p13** was obtained (99 mg, 81%) and used in the next step without further purification. LC $t_R$ = 4.76 min, MS (ESI) *m/z:* 439.39 $[M+H]^+$. $^1H$ NMR (500 MHz, DMSO-$d_6$) δ 10.15 (br s, 1H), 7.48 – 7.46 (m, 1H), 7.46 (d, *J* = 8.5 Hz, 1H), 7.12 (d, *J* = 8.5 Hz, 2H), 7.05 (d, *J* = 8.5 Hz, 2H), 6.84 (d, *J* = 8.7 Hz, 2H), 4.10 – 4.05 (m, 2H), 3.78 (s, 2H), 3.73 – 3.68 (m, 4H), 3.56 – 3.52 (m, 1H), 3.05 – 3.00 (m, 4H), 1.76 – 1.70 (m, 1H), 1.67 – 1.60 (m, 2H), 1.53 – 1.43 (m, 2H), 1.16 – 1.13 (m, 3H), 0.90 – 0.85 (m, 9H) $^{13}C$ NMR (126 MHz, DMSO-$d_6$)[1] δ 169.84, 166.86, 149.36, 137.17, 136.70, 132.16, 129.12 (2C), 128.85 (2C), 119.39 (2C), 115.34 (2C), 66.12 (2C), 60.61, 50.89, 48.75 (2C), 37.43, 25.64, 22.68, 21.94, 14.04

[1] Overlapping signals not possible to accurately integrate

*N*1-hydroxy-2-isobutyl-*N*3-(4-(4-morpholinobenzyl)phenyl)malonamide (**3a1**)

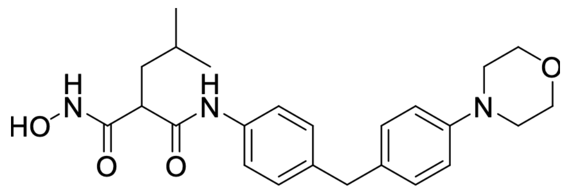

Compound **3a1** was synthesized according to General Procedure C using compound **p13** (50 mg, 0.114 mmol), $NH_2OH$ (50% in water, 230 µL, 3.76 mmol), and KCN (1.48 mg, 0.022 mmol) in a 1:1 mixture of THF/MeOH (0.57 mL). The reaction mixture was stirred for 72 h. The crude product was purified by flash chromatography on silica gel followed by preparative HPLC ($H_2O$ + 0.05% FA/ ACN + 0.05% FA, 95:5 to 5:95) to afford compound **3a1** as a white solid (33 mg, 68%). LC $t_R$ = 3.78 min, MS (ESI) *m/z:* 425.26 $[M+H]^+$. $^1H$ NMR (500 MHz, DMSO-$d_6$) δ 10.52 (br s, 1H), 9.68 (br s, 1H), 8.95 (br s, 1H), 7.46 (d, *J* = 8.4 Hz, 2H), 7.11 (d, *J* = 8.5 Hz, 2H), 7.05 (d, *J* = 8.7 Hz, 2H), 6.84 (d, *J* = 8.7 Hz, 2H), 3.77 (s, 2H), 3.75–3.67 (m, 4H), 3.19 (t, *J* = 7.6 Hz, 1H), 3.07 – 2.99 (m, 4H), 1.70 – 1.64 (m, 2H), 1.51–1.42 (m, 1H), 0.87 (dd, *J* = 6.6, 3.6 Hz, 7H). $^{13}C$ NMR (126 MHz, DMSO-$d_6$) δ 168.02, 166.86, 149.82, 137.44, 137.12, 132.62, 129.60 (2C), 129.24 (2C), 119.95 (2C), 115.79 (2C), 66.59 (2C), 50.34, 49.21 (2C), 38.56, 26.17, 22.88, 22.72.

4-Methyl-2-((4-(4-morpholinobenzyl)phenyl)carbamoyl)pentanoic acid (**p14**)

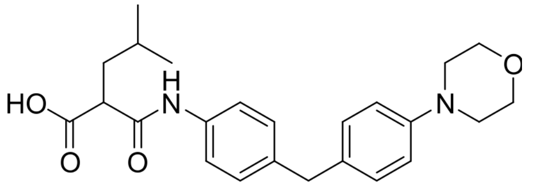


Compound **p14** was synthesized according to General Procedure D using ester **p13** (100 mg, 0.228 mmol) in THF (1 mL) and aqueous LiOH (2 M, 1 mL). The reaction mixture was stirred for 1.5 h. Compound **p14** was obtained and used in the next step without further purification (93 mg, quantitative). LC $t_R$ = 4.25 min, MS (ESI) *m/z:* 411.33 $[M+H]^+$. $^1H$ NMR (500 MHz, DMSO-$d_6$) δ 12.64 (br s, 1H), 10.12 (s, 1H), 7.48 (d, *J* = 8.4 Hz, 2H), 7.12 (d, *J* = 8.4 Hz, 2H), 7.05 (d, *J* = 8.5 Hz, 2H), 6.85 (d, *J* = 8.5 Hz, 2H), 3.77 (s, 2H), 3.73 – 3.68 (m, 4H), 3.47–3.44 (m, 2H), 3.06–2.98 (m, 4H), 1.75–1.68 (m, 1H), 1.62–1.58 (m, 2H), 1.54–1.43 (m, 2H), 0.89–0.85 (m, 9H). $^{13}C$ NMR (126 MHz, DMSO-$d_6$) δ 171.50, 167.46, 149.39, 137.02, 136.91, 132.23, 129.16 (2C), 128.89 (2C), 119.28 (2C), 115.36 (2C), 66.16 (2C), 50.99, 48.78 (2C), 37.41, 25.70, 22.84, 21.97.

*N*1-ethoxy-2-isobutyl-*N*3-(4-(4-morpholinobenzyl)phenyl)malonamide (**3a**)

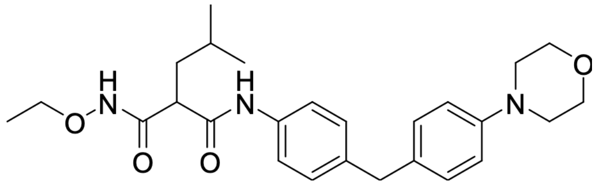


Compound **3a** was synthesized according to General Procedure A using carboxylic acid **p14** (120 mg, 0.292 mmol), *O*-ethylhydroxylamine hydrochloride (34 mg, 0.350 mmol), T3P (50% in EtOAc, 258 µL, 0.876 mmol), and $Et_3N$ (122 µL, 0.876 mmol) in $CH_2Cl_2$ (1.46 mL). The reaction mixture was stirred for 16 h. The crude product was purified by flash chromatography on silica gel to afford compound **3a** as a yellow oil (158 mg, 90%). LC $t_R$

= 4.26 min, MS (ESI) *m/z:* 454.42 $[M + H]^+$. $^1$H NMR (500 MHz, DMSO-$d_6$) δ 10.96 (s, 1H), 9.70 (s, 1H), 7.46 (d, *J* = 8.5 Hz, 2H), 7.12 (d, *J* = 8.4 Hz, 2H), 7.05 (d, *J* = 8.5 Hz, 2H), 6.84 (d, *J* = 8.7 Hz, 2H), 3.80 – 3.76 (m, 4H), 3.75 – 3.67 (m, 4H), 3.19 (t, *J* = 7.6 Hz, 1H), 3.06 – 3.00 (m, 4H), 1.72 – 1.61 (m, 2H), 1.53 – 1.44 (m, 1H), 1.12 (t, *J* = 7.0 Hz, 3H), 0.87 (br dd, *J* = 6.6, 1.8 Hz, 6H). $^{13}$C NMR (126 MHz, DMSO-$d_6$) δ 167.31, 166.25, 149.35, 137.06, 136.64, 132.15, 129.12 (2C), 128.78 (2C), 119.54 (2C), 115.32 (2C), 70.55, 66.12 (2C), 50.05, 48.75 (2C), 37.89, 25.78, 22.34, 22.24, 22.22, 13.37.
